# Temporal Dynamics of Genetically Heterogeneous Extended-Spectrum Cephalosporin Resistant *Escherichia coli* Bloodstream Infections

**DOI:** 10.1101/2023.02.07.527510

**Authors:** William C Shropshire, Benjamin Strope, Selvalakshmi Selvaraj Anand, Jordan Bremer, Patrick McDaneld, Micah M Bhatti, Anthony R Flores, Awdhesh Kalia, Samuel A Shelburne

## Abstract

Extended-spectrum cephalosporin resistant *Escherichia coli* (ESC-R-*Ec*) is an urgent public health threat with sequence type clonal complex 131 (STc131), phylogroup B2 strains being particularly concerning as the dominant cause of ESC-R-*Ec* infections. To address the paucity of recent ESC-R-*Ec* molecular epidemiology data in the United States, we used whole genome sequencing (WGS) to fully characterize a large cohort of invasive ESC-R-*Ec* at a tertiary care cancer center in Houston, Texas collected from 2016-2020. During the study timeframe, there were 1154 index *E. coli* bloodstream infections (BSIs) of which 389 (33.7%) were ESC-R-*Ec*. Using time series analyses, we identified a temporal dynamic of ESC-R-*Ec* distinct from ESC-susceptible *E. coli* (ESC-S-*Ec*), with cases peaking in the last six months of the calendar year. WGS of 297 ESC-R-*Ec* strains revealed that while STc131 strains accounted for ∼45% of total BSIs, the proportion of STc131 strains remained stable across the study time frame with infection peaks driven by genetically heterogeneous ESC-R-*Ec* clonal complexes. *Bla*_CTX-M_ variants accounted for most β-lactamases conferring the ESC-R phenotype (89%; 220/248 index ESC-R*-Ec*), and amplification of *bla*_CTX-M_ genes was widely detected in ESC-R-*Ec* strains, particularly in carbapenem non-susceptible, recurrent BSI strains. *Bla*_CTX-M-55_ was significantly enriched within phylogroup A strains, and we identified *bla*_CTX-M-55_ plasmid-to-chromosome transmission occurring across non-B2 strains. Our data provide important information regarding the current molecular epidemiology of invasive ESC-R-*Ec* infections at a large tertiary care cancer center and provide novel insights into the genetic basis of observed temporal variability for these clinically important pathogens.

**IMPORTANCE:** Given that *E. coli* is the leading cause of worldwide ESC-R *Enterobacterales* infections, we sought to assess the current molecular epidemiology of ESC-R-*Ec* using a WGS analysis of many BSIs over a five-year period. We identified fluctuating temporal dynamics of ESC-R-*Ec* infections, which has also recently been identified in other geographical regions such as Israel. Our WGS data allowed us to visualize the stable nature of STc131 over the study period and demonstrate a limited, but genetically diverse group of ESC-R-*Ec* clonal complexes are detected during infection peaks. Additionally, we provide a widespread assessment of β-lactamase gene copy number in ESC-R-*Ec* infections and delineate mechanisms by which such amplifications are achieved in a diverse array of ESC-R-*Ec* strains. These data suggest that serious ESC-R-*Ec* infections are driven by a diverse array of strains in our cohort and impacted by environmental factors suggesting that community-based monitoring could inform novel preventative measures.

## INTRODUCTION

Extended-spectrum β-lactamase (ESBL) producing *Enterobacterales* (ESBL-E) were ranked as a serious antimicrobial resistance (AMR) threat by the Centers for Disease Control and Prevention in 2019 (1). A recent study of multi-drug resistant (MDR) infections in hospitalized patients indicated a 53.3% increased incidence rate in estimated ESBL-E cases from 2012 to 2017 in the United States (US) (2). Other sources of US data have indicated similar trends, such as a 138% increase in resistance to 3^rd^ generation cephalosporins amongst invasive *E. coli* strains from 2009 to 2016 (3, 4). Furthermore, multiple population-based studies indicate that ESBL-E bloodstream infections may have a higher likelihood of adverse outcomes compared to non-ESBL-E bloodstream infections (5–7).

The key impact of ESBL enzymes is 3^rd^ generation cephalosporin resistance (*i.e.,* extended-spectrum cephalosporin resistance; ESC-R). Although ESBL enzymes account for the vast majority of the associated ESC-R phenotype, there are other mechanisms in *Enterobacterales*, such as plasmid-borne AmpC enzymes (8). Therefore, broadening our scope of potential ESC-R mechanisms provides a more holistic approach to understanding transmission dynamics of these ESC-R genes (9). Amongst ESC-R *Enterobacterales*, some 50% or greater of infections are due to *Escherichia coli* (10, 11). ESC-R *E. coli* (ESC-R-*Ec*) in the 1980s and 1990s primarily harbored ESBL variants of TEM and SHV enzymes (12); however, this changed in the early 2000s with the increasing spread of CTX-M enzymes (13). The rise of CTX-M enzyme producing *E. coli* coincided with a marked increase in sequence type (ST) 131 strains (14). In particular, fluoroquinolone resistant sub-populations of ST131 (*i.e.,* ST131-*H*30 subclade C) strongly associated with *bla*_CTX-M-15_ and *bla*_CTX-M-27_ carriage, have been shown to be the predominant cause of ESC-R-*Ec* infections in the US (15). However, whether the increasing number of ESC-R-*Ec* infections in the US is driven by further expansion of ST131 strains is not well understood.

Application of WGS to large cohorts of temporally diverse *E. coli*, including ESBL producing isolates, from local and nationwide surveys has revealed common as well as unique molecular epidemiology features (16–23). Consistently, *E. coli* strains causing human infections are pauci-clonal with domination of a small number of STs relative to the overall population (16–18, 24). Many analyses indicate ST131 rose to become the leading *E. coli* ST isolated from humans by 2010 (19, 25). However, studies in England and Calgary have found that ST131 has remained at a relatively steady proportion of overall *E. coli* infections over the past five to ten years (16, 19, 22) whereas investigations in France, Spain, and Norway have identified a continued increase in emergent ST131 subclades (17, 20, 21). Geographical variation in relative ST131 proportion changes may reflect differential ST131 cladal adaptation within ecological niches, with advantageous traits failing to fixate in populations due to negative frequency-dependent selection, reflected by the eventual stability of ST131 within the broader *E. coli* population (26).

A comprehensive US study found that ST131 accounted for 50% of all ESC-R-*Ec* strains in 2012 with an estimated increase of 50% from 2007 to 2011 in the percentage of ST131 *E. coli* (27). Given the lack of recent longitudinal data analyzing ESC-R-*Ec* in the US, we sought to gain insight into the molecular epidemiology of ESC-R-*Ec* by analyzing bloodstream infection causing strains at our institution from 2016-2020.

## RESULTS

### Observed peaks in ESC-R-*Ec* BSIs occurred temporally within our sampling frame

We identified total *Escherichia coli* bloodstream infections (*Ec*-BSI) at our institution that occurred from March 1^st^, 2016, to December 31^st^, 2020. There were 1370 total *Ec*-BSIs with 84.2% (1154/1370) being index *Ec*-BSI cases and 15.8% (216/1370) being recurrent (see methods for definitions). The overall ESC-R-*Ec* proportion was 33.7% (389/1154) among index *Ec*-BSI cases. The proportion of ESC-R-*Ec* infections that were recurrent (121/510, 23.7%) was significantly higher compared to recurrent extended-spectrum cephalosporin susceptible *E. coli* (ESC-S-*Ec*) cases (95/860, 11%, χ^2^ *P*-value < 0.001).

The non-normalized temporal trends of ESC-R-*Ec* within this 4+ year timeframe is presented in Table S1. Given that changes in absolute yearly ESC-R-*Ec* might be reflective of variance in the total population seen at our institution, we utilized monthly admissions as a surrogate of the hospital population to calculate prevalence rates. We found no statistically significant linear increase in ESC-R-*Ec* rates over the course of the study (linear regression one-sample t-test *P*-value=0.20). Notably, when calculating *Ec*-BSI prevalence rates stratified by ESC status, we identified an oscillating pattern of total *Ec*-BSI prevalence rates corresponding to peaks and valleys (dotted grey loess curve; **Figure 1**) that primarily correlated with ESC-R-*Ec* prevalence rates (solid red loess curve; **Figure 1**) rather than ESC-S-*Ec* prevalence rates (solid blue curve, **Figure 1**). Stratifying the sampling frame based on these observed increased infection peaks (**Figure 1; Table 1**), ESC-R-*Ec* prevalence rates were greatest during window 2 (1.89 BSI cases/1000 admissions), window 4 (2.09 BSI cases/1000 admissions), and window 6 (2.24BSI cases/1000 admissions). The mean peak ESC-R-*Ec* prevalence rate (2.07 BSI cases/1000 admissions) was 60% greater than the average of 1.29 BSI cases/1000 admissions during non-peak periods (P < 0.001 for monthly ESC-R-*Ec* rates during peak vs. non-peak windows by Student’s t-test).

**Figure 1.**
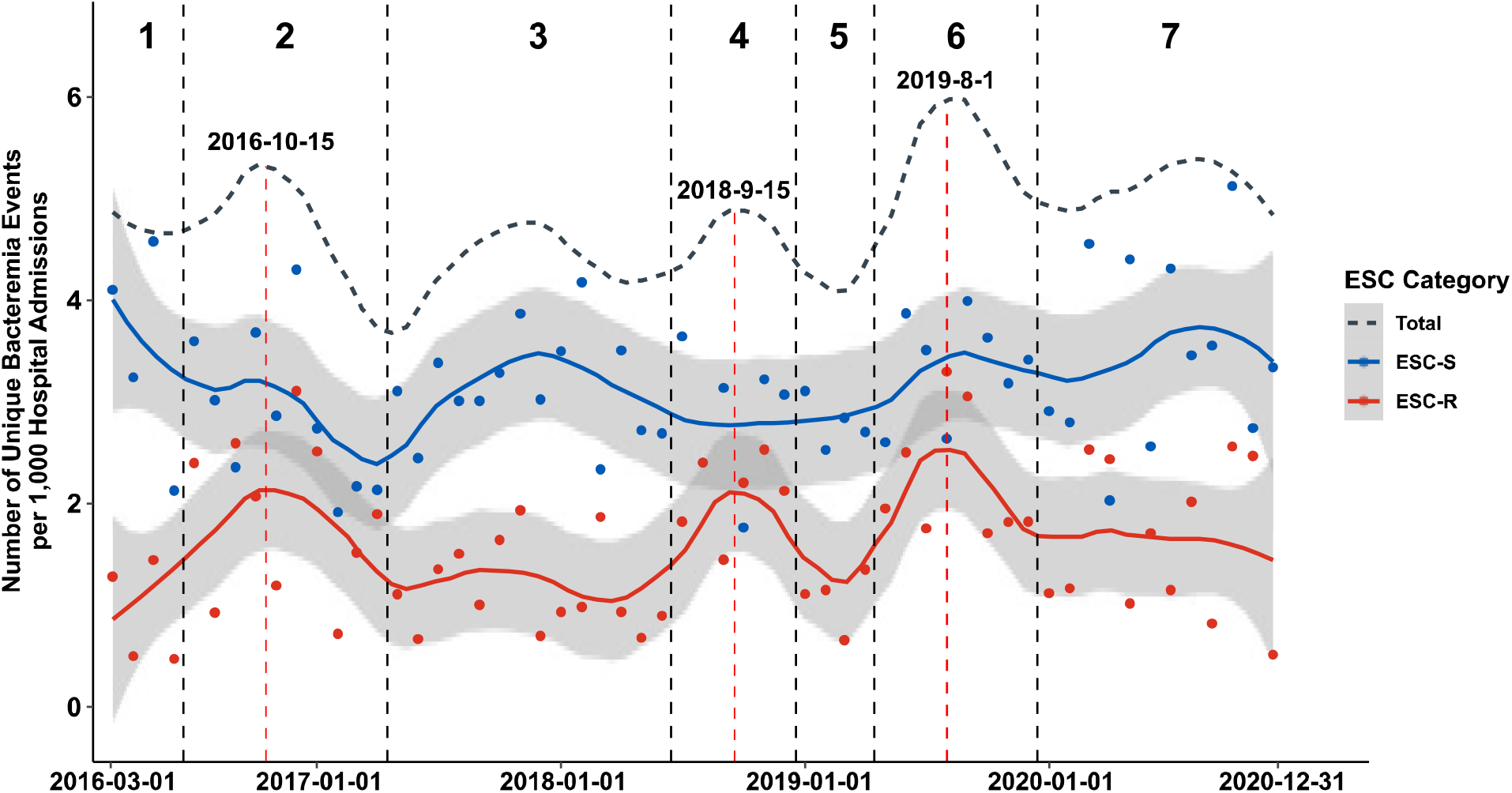
Number of Bacteremia Events Observed per Month from March 2016 to December 2020 at MD Anderson Cancer Center. Number of index *E. coli* bacteremia events per 1000 hospital admissions for both ESC-R (red) and ESC-S (blue) isolates. A local regression curve for each respective ESC category is provided with 95% confidence intervals (gray area). The dotted gray line represents the prevalence trend for total *E. coli* bacteremia events. Windows of peak infection estimates were observed in windows 2, 4, and 6 as demarcated above by vertical dotted black lines with estimated peak date indicated within peak regions (vertical dotted red lines).

**Table 1:**
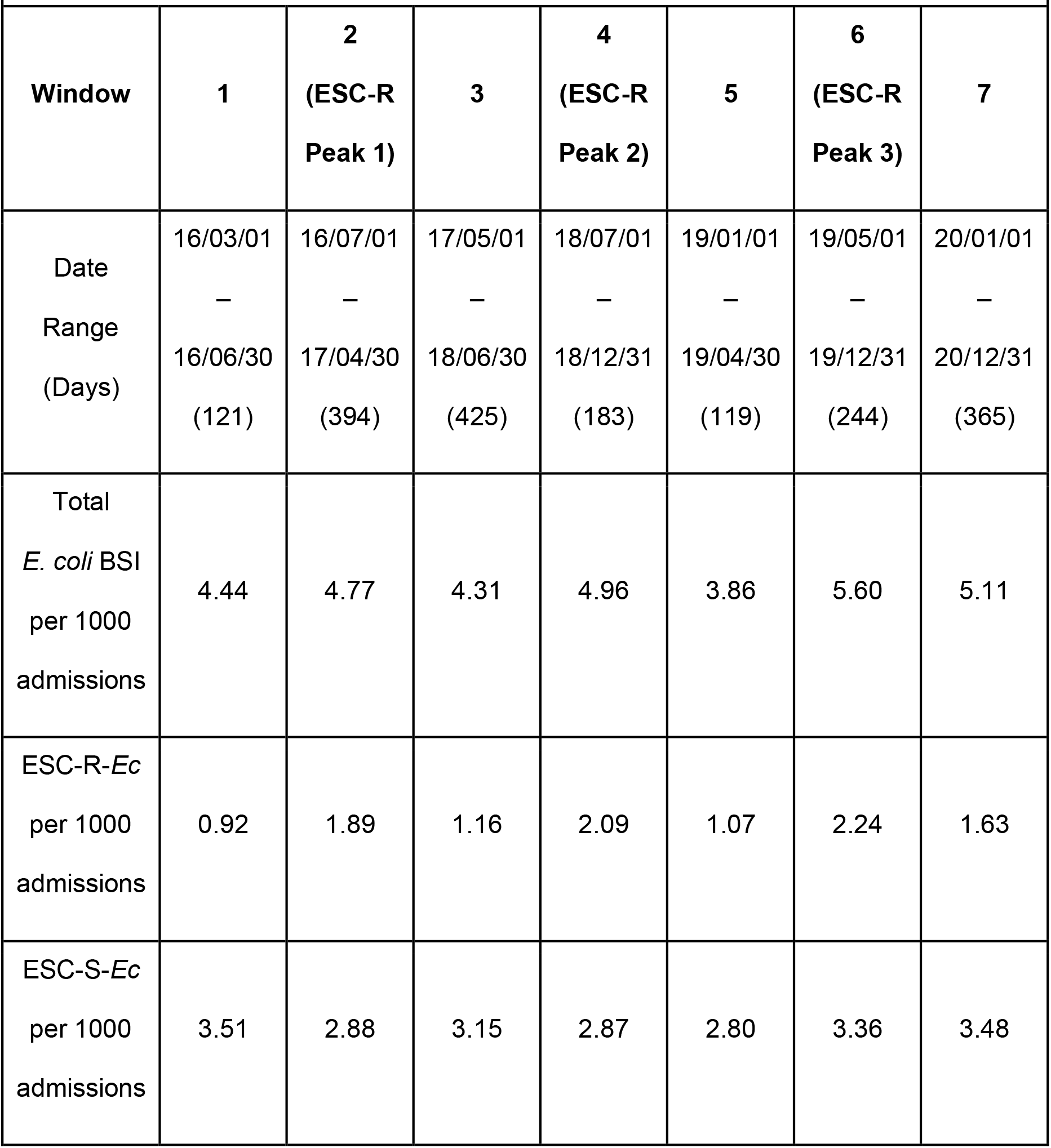
Windows of *E. coli* Bloodstream Infection Rates per 1000 Hospital Admissions from 2016-03-01 to 2020-12-31.

We performed time series analyses and found that while both ESC-R-*Ec* and ESC-S-*Ec* had non-significant monotonic trends, there was evidence of a non-monotonic trend for ESC-R-*Ec* that occurred up to the start of the COVID-19 pandemic in March 2020 (WAVK test-statistic =2.7; *P*-value = 0.016), which was not evident in ESC-S-*Ec* (WAVK test-statistic =0.2; *P*-value = 0.877). To determine whether ESC-R-*Ec* and ESC-S-*Ec* prevalence rates were distinct, we compared pairwise month-to-month prevalence and found no statistically significant correlation (R=0.14, Pearson *P*-value = 0.31; Fig. S1A) suggesting independent temporal trends. Given the periodic trends noted in **Figure 1**, we assessed whether variation in *Ec-*BSIs corresponded to the time of year by dividing the index infections into the 1^st^ or 2^nd^ half of the year. We found a significantly higher average of ESC-R-*Ec* (7.9 ± 0.5 vs. 5.5 ± 0.6 BSIs/month; *P*-value = 0.003) but not ESC-S-*Ec* (13.8 ± 0.5 vs. 12.5 ± 0.6; *P*-value =0.09) occurring in the last 6 months relative to the initial 6 months of each year (Fig. S1B) respectively. Taken together, these data trends suggest temporal dynamics in ESC-R-*Ec* independent of ESC-S-*Ec* strains with a potential seasonal component.

### Peaks in ESC-R *Ec*-BSIs are composed of genetically heterogeneous strains from multiple sequence type complexes

From the 389 index and 121 recurrent ESC-R-*Ec* strains identified during the study period, we generated assemblies for 297 ESC-R-*Ec* isolates including 64% (248/389) of index and 41% (49/121) of recurrent ESC-R-*Ec* strains. The quality of assemblies is shown in Table S2. In total there were 7 phylogroups detected across the full sampling frame (**Table 2**). The majority of index ESC-R-*Ec* belonged to phylogroup B2 (50.8%; 126/248), followed by A (19.8%; 49/248), D (12.1%; 30/248), F (7.7%; 19/248), B1 (6.5%; 16/248), C (2.8%; 7/248), and G (0.4%; 1/248).

**Table 2:**
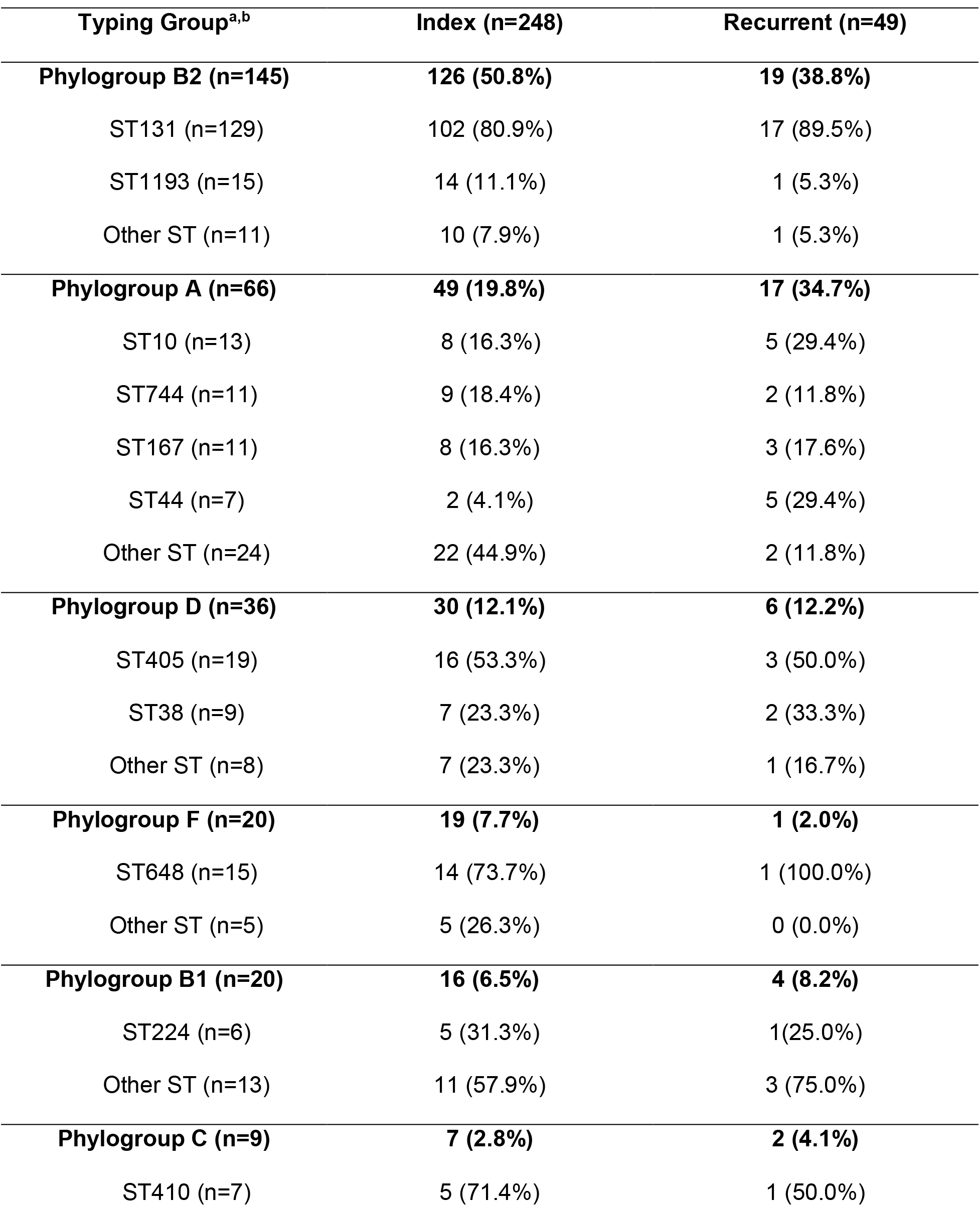
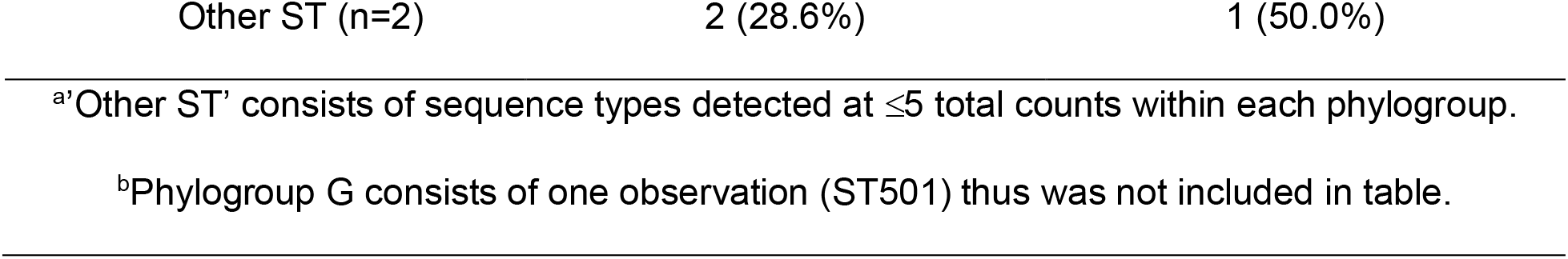
Overview of Sequence Type Distribution Across Each Phylogroup from Sequenced Index and Recurrent ESC-R *E.* coli Isolates.

| Typing Group<sup>a,b</sup> | Index (n=248) | Recurrent (n=49) |
| --- | --- | --- |
| **Phylogroup B2 (n=145)** | **126 (50.8%)** | **19 (38.8%)** |
| ST131 (n=129) | 102 (80.9%) | 17 (89.5%) |
| ST1193 (n=15) | 14 (11.1%) | 1 (5.3%) |
| Other ST (n=11) | 10 (7.9%) | 1 (5.3%) |
| **Phylogroup A (n=66)** | **49 (19.8%)** | **17 (34.7%)** |
| ST10 (n=13) | 8 (16.3%) | 5 (29.4%) |
| ST744 (n=11) | 9 (18.4%) | 2 (11.8%) |
| ST167 (n=11) | 8 (16.3%) | 3 (17.6%) |
| ST44 (n=7) | 2 (4.1%) | 5 (29.4%) |
| Other ST (n=24) | 22 (44.9%) | 2 (11.8%) |
| **Phylogroup D (n=36)** | **30 (12.1%)** | **6 (12.2%)** |
| ST405 (n=19) | 16 (53.3%) | 3 (50.0%) |
| ST38 (n=9) | 7 (23.3%) | 2 (33.3%) |
| Other ST (n=8) | 7 (23.3%) | 1 (16.7%) |
| **Phylogroup F (n=20)** | **19 (7.7%)** | **1 (2.0%)** |
| ST648 (n=15) | 14 (73.7%) | 1 (100.0%) |
| Other ST (n=5) | 5 (26.3%) | 0 (0.0%) |
| **Phylogroup B1 (n=20)** | **16 (6.5%)** | **4 (8.2%)** |
| ST224 (n=6) | 5 (31.3%) | 1 (25.0%) |
| Other ST (n=13) | 11 (57.9%) | 3 (75.0%) |
| **Phylogroup C (n=9)** | **7 (2.8%)** | **2 (4.1%)** |
| ST410 (n=7) | 5 (71.4%) | 1 (50.0%) |

<sup>a</sup>'Other ST' consists of sequence types detected at $\leq 5$ total counts within each phylogroup.
<sup>b</sup>Phylogroup G consists of one observation (ST501) thus was not included in table.

We identified a similar distribution for recurrent ESC-R-*Ec* strains with B2 (38.8%; 19/49) as the most common phylogroup followed by A (34.7%; 17/49), D (12.2%; 6/49), B1 (8.2%; 4/49), C (4.1%; 2/49), and F (2.0%; 1/49). In terms of sequence types (STs) (**Table 2**), ST131 was the most common for both index (n = 102, 41%) and recurrent strains (n = 17, 35%). A further breakdown of each of the most common STs detected by phylogroup can be found in **Table 2**.

To investigate the molecular epidemiology underlying the temporal dynamics of our ESC-R-*Ec* population, we analyzed isolates characterized by phylogroup across infection windows (**Figure 2**). **Figure 2A-B** provides an illustration of the phylogroup distribution throughout the infection windows which remained remarkably stable over time with no statistically significant trends (Fisher’s exact test simulated *P*-value= 0.95). There were sporadic increases in the absolute numbers of various phylogroups during the peak infection periods such as phylogroup A during windows 2 and 6, as well as phylogroup B1 and phylogroup F during window 4. We collapsed the “non-peak” (*i.e.,* windows 1, 3, 5, and 7) and “peak” (*i.e.,* windows 2, 4, and 6) periods together and again found no statistically significant differences in phylogroup proportions between peak and non-peak periods (Fisher’s exact test *P*-value= 0.42). The dominant B2 phylogroup accounted for 58% of infections during the non-peak windows and 47% during the peak windows (Fisher’s exact test *P*-value= 0.15).

**Figure 2.**
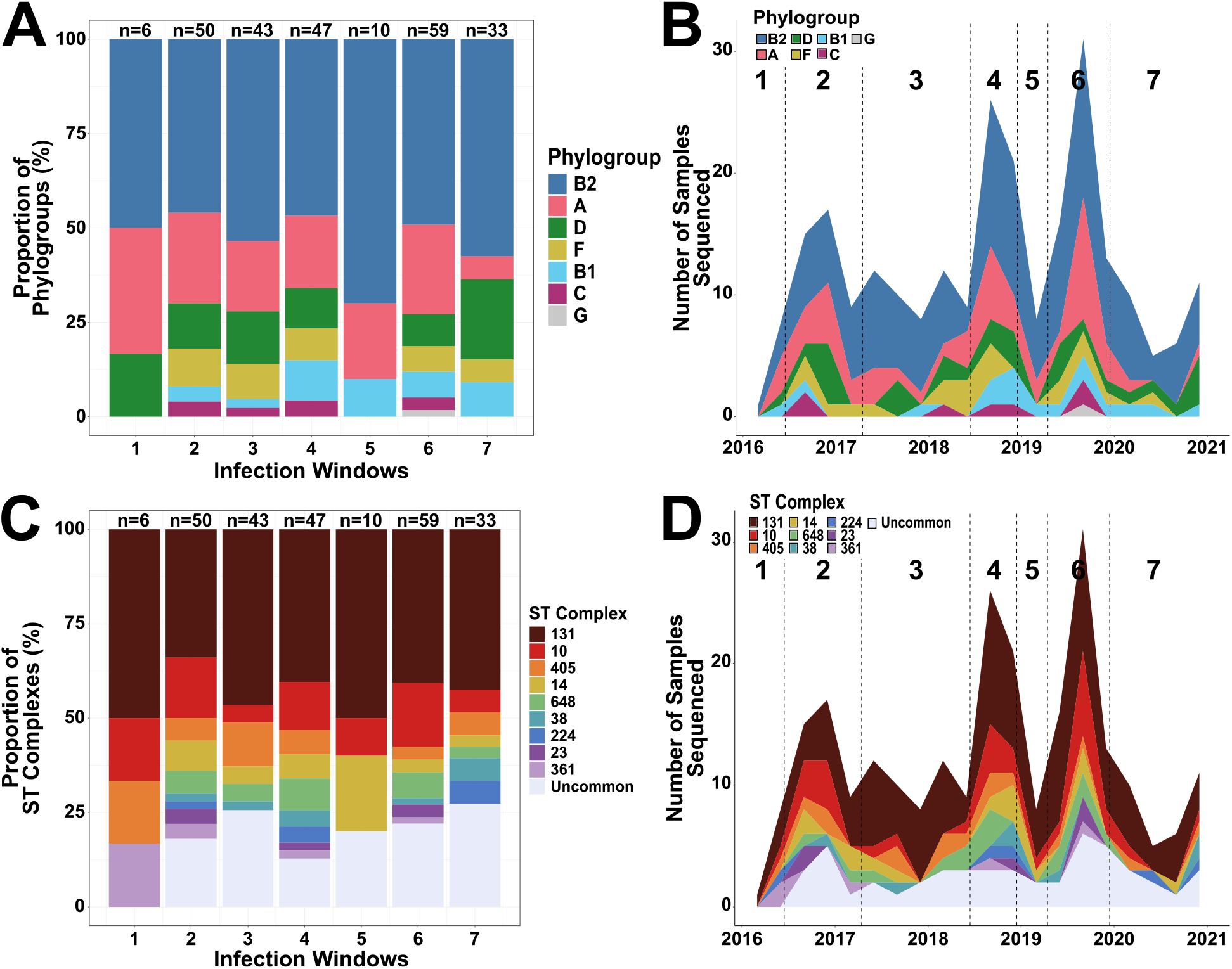
Temporal Distribution of ESC-R *Ec*-BSI Isolates. The proportional distribution and absolute frequency counts by phylogroup **(A-B)** and clonal complex **(C-D)** respectively across each infection window. **(A-C)** Group totals are listed above each of the windows. **(B-D)** Data shown are line graphs connecting month-to-month frequency counts colored by proportions of phylogroup **(B)** and clonal complex **(D)** as indicated in the legend.

We grouped index strains by sequence type clonal complexes (*i.e.,* single locus ST variants within a 7-housekeeping gene schema) to further analyze temporal trends. The leading sequence type complexes (STc) were STc131 (n = 109, 44%), STc10 (n = 35, 14%), STc405 (n = 16, 7%), STc14 (n = 14, 6%), STc648 (n = 14 (6%), and STc38 (n = 10, 4%). For testing purposes, we collapsed STcs with less than 5 strains into an “uncommon” category (n = 32, 13%). Similar to our phylogroup analysis, we did not identify any significant trends for a particular STc over the study period (Fisher’s exact test simulated *P*-value= 0.91). However, there were temporary absolute increases in specific STc infections within peak infection windows such as “uncommon” STcs within window 2, STc14 and STc648 within window 4, and STc10 within window 6 (**Figure 2C-D**). Importantly, STc131 remained stable in prevalence across each of the infection windows (χ^2^ *P*-value = 0.9) with STc131 strains accounting for 50% of infections during non-peak windows compared to 40% of infections during peak infection windows (Fisher’s exact test *P*-value= 0.15). Temporal information regarding isolates sequenced per window including the proportion of isolates that were STc131 is provided in Table S3. There were no peak infection periods in which STc131 strains accounted for a higher percentage of infections relative to non-peak periods (Table S3). Similarly, there were no statistically significant proportion differences across windows in ST1193, the other most commonly prevalent B2 isolates (Fisher’s exact test *P*-value= 0.48) nor in the second most common, STc10 (Fisher’s exact test *P*-value= 0.74). Thus, these data indicate that peaks of ESC-R-*Ec* observed temporally over our study period were driven by a diverse combination of ESC-R-*Ec* isolates rather than expansion of STc131 strains.

**Table 3:** Copy Number Variants of β-lactamase Encoding Genes.

|  | CTX-M-15 | CTX-M-27 | CTX-M-55 | CTX-M-14 | Other<br>CTX-M |
| --- | --- | --- | --- | --- | --- |
| **Total** |  |  |  |  |  |
| **Count** | 161 | 37 | 37 | 23 | 10 |
| **Mean (SD)** | 1.62 (1.29) | 1.87 (1.89) | 2.50 (2.65) | 2.61 (2.57) | 3.55 (3.35) |
| **Median (IQR)** | 1.11 (1.19) | 1.40 (0.80) | 1.50 (1.43) | 1.42 (2.21) | 1.31 (5.36) |
| **Range** | 0.37-8.72 | 0.42-9.10 | 0.62-12.9 | 0.64-10.8 | 0.60-9.45 |
| **Phylogroup B2** |  |  |  |  |  |
| **Count** | 92 | 33 | 1 | 6 | 3 |
| **Mean (SD)** | 1.41 (0.96) | 1.12 (0.36) | 3.41 | 2.53 (2.06) | 4.80 (3.66) |
| **Median (IQR)** | 1.03 (0.54) | 1.20 (0.354) | NA | 1.75 (1.83) | 6.54 (3.33) |
| **Range** | 0.44-5.84 | 0.42-9.10 | NA | 0.88-6.33 | 0.60-7.26 |
| **Phylogroup A** |  |  |  |  |  |
| **Count** | 31 | 3 | 19 | 5 | 2 |
| **Mean (SD)** | 1.66 (1.72) | 1.12 (0.36) | 2.39 (2.09) | 1.65 (1.42) | 1.08 (0.13) |
| **Median (IQR)** | 1.07 (0.97) | 1.20 (0.35) | 1.68 (1.82) | 1.01 (0.31) | 1.08 (0.09) |
| **Range** | 0.50 - 8.72 | 0.72 - 1.44 | 0.62 - 7.20 | 0.78-4.16 | 0.98-1.17 |
| **Phylogroup D** |  |  |  |  |  |
| **Count** | 21 | 1 | 5 | 4 | 3 |
| **Mean (SD)** | 2.02 (1.48) | 1.8 | 1.25 (0.49) | 1.80 (0.98) | 3.90 (4.81) |
| **Median (IQR)** | 2.00 (1.62) | NA | 1.30 (0.53) | 1.43 (0.54) | 1.31 (4.25) |
| **Range** | 0.37-5.14 | NA | 1.00-1.96 | 1.10-3.26 | 0.94-9.45 |

SD = Standard Deviation; IQR = Interquartile Range; Copy Number Variants are estimated based on CONVICT results

### *Bla*_CTX-M_ variants are dominant encoding genes conferring inferred ESC-R phenotype

We searched the WGS data for genes potentially conferring the ESC-R phenotype, which we defined as those encoding ESBLs, *i.e.*, TEM, SHV, CTX-M encoding genes with a 2be Jacoby-Bush classification (12), as well as CMY and carbapenemase encoding genes. Of the 248 index isolates, 236 (95%) had at least one gene expected to confer the ESC-R phenotype with CTX-M variants being present in 220 strains (89%) and CMY variants found in 21 strains (8.4%) (*N.b.,* strains could contain multiple ESC-R encoding genes). A single ST131 strain carried an ESBL *bla*_TEM-19_, two strains carried ESBL *bla*_SHV-12_, one ST361 strain carried *bla*_OXA-232_, and three STc10 strains harbored *bla*_NDM-5_. The most common CTX-M encoding gene variants detected within index ESC-R-*Ec* isolates were *bla*_CTX-M-15_ (n=134), *bla*_CTX-M-27_ (n=29), *bla*_CTX-M-55_ (n=26), *bla*_CTX-M-14_ (n=19), and multiple *bla*_CTX-M_ variant carriage (n=7). A similar distribution was observed for recurrent ESC-R-*Ec* isolates: *bla*_CTX-M-15_ (n=26), *bla*_CTX-M-27_ (n=7), *bla*_CTX-M-55_ (n=8), and *bla*_CTX-M-14_ (n=3). Of the 12 index strains that did not carry a known ESC-R phenotype determinant, six contained *bla*_OXA-1_, five encoded non-ESBL TEM enzymes, and only one isolate lacked a known exogenous β-lactamase encoding gene. The preponderance of CTX-M with a smaller role of CMY in conferring the ESC-R phenotype in *E. coli* is consistent with previously published results in the US (9, 10).

### Temporal whole genome analysis indicates low level of clonality amongst ESC-R-*Ec*

To gain further insights into the molecular epidemiology driving the temporal dynamics of ESC-R-*Ec*, we used WGS data to analyze the population structure for all 297 isolates. The core gene inferred maximum likelihood phylogeny based on 2,695 core genes (*i.e.,* genes present in ≥99% of the 297 ESC-R *Ec*-BSI Isolates) is shown in **Figure 3**. Consistent with previous WGS studies of ESC-R *E. coli* population structure (9, 18), we observed significant variation in the degree of genetic similarity amongst strains of distinct STcs. For example, there was only an average of 28 core gene SNPs separating a given STc14 strain from its nearest neighbor compared to 1518 for STc10 strains. Similarly, we identified several previously described subclades within the broader STc131 population (15, 20, 28). By overlying infection window with our WGS phylogeny, we sought to determine whether the increased resolution would facilitate detection of highly related strains driving infection peaks. However, in line with our phylogroup and STc-based analyses, we did not observe large numbers of phylogenetically clustered index strains occurring during infection windows (**Figure 3**). Consistent with the notion of similar genetic diversity being present during peak and non-peak infection windows, there was no difference in pairwise core gene SNP distance between one isolate and its most closely related strain during the various windows (Kruskal-Wallis test *P*-Value=0.81; Fig S2).

**Figure 3.**
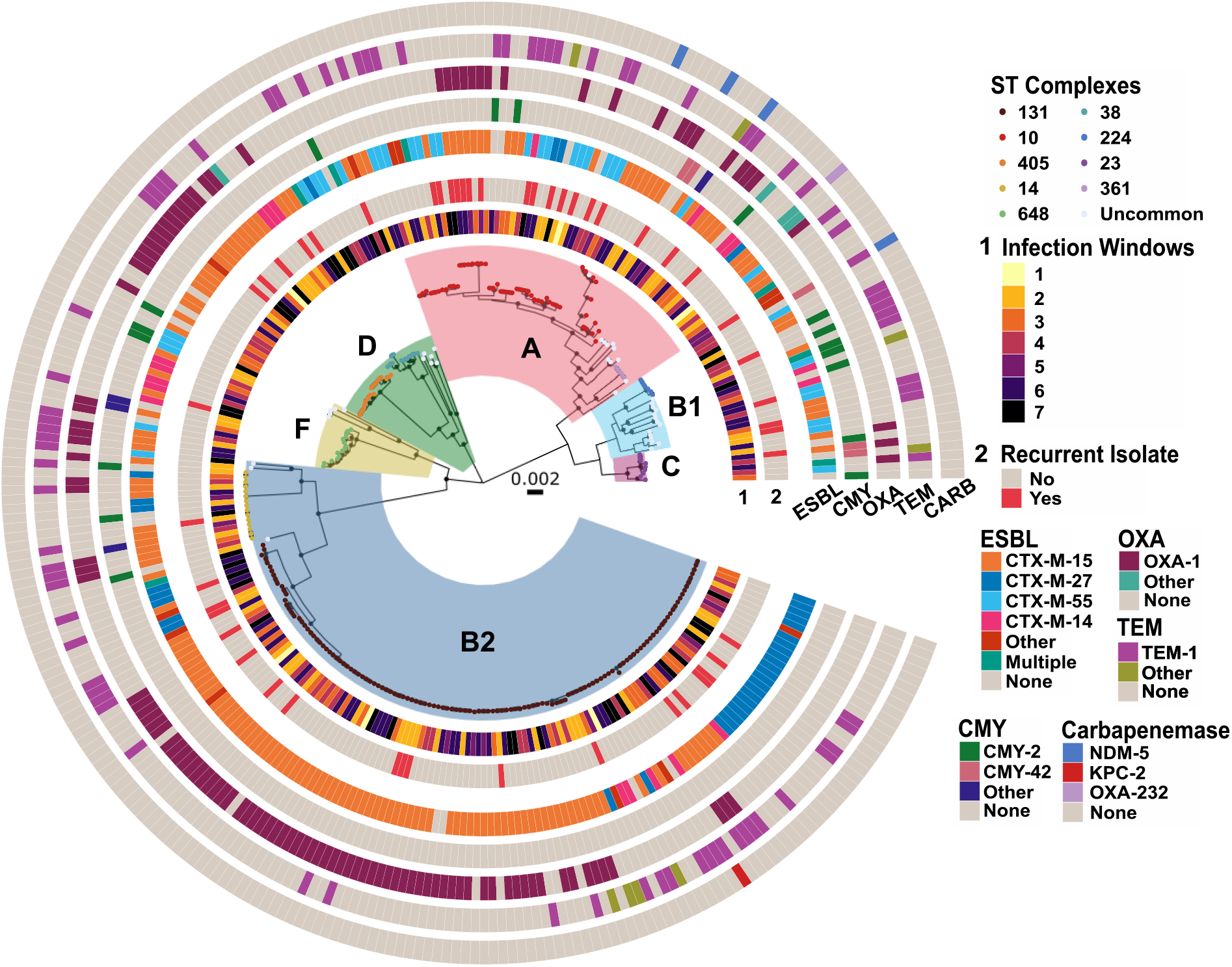
Population Structure of ESC-R *Ec*-BSI isolates. A core gene alignment inferred maximum likelihood phylogenetic tree with phylogroups highlighted within each monophyletic group. Branch tip colors indicate clonal complex number as shown in the figure inset with clonal complexes observed <5 counts grouped as “Uncommon”. Rings corresponding to infection window (1), recurrence (2), and β-lactamases are labelled respectively. Internal nodes labelled with black circles have SH-aLRT ≥ 80% and UFboot ≥ 95%.

To search for clusters of closely genetically related strains both within peak infection windows and throughout the study period, we measured the pairwise SNP distances of index isolates using recombination free core genome alignments from the 12 most commonly observed STs in our cohort, which make up 79.8% of the total population (198/248). We identified seven groups of two or more isolates (*i.e.*, genetic clusters) from unique patients with <10 core genome pairwise SNPs (Fig S3). The maximum number of closely related isolates within a peak infection window was two strains whereas the maximum cluster size was three strains, with duration between isolates from the same cluster ranging from 5 days to 777 days (Fig. S3). These data indicate that even at the WGS level, peaks of ESC-R-*Ec* are driven by genetically heterogenous isolates with few signals indicating clonal outbreaks across each infection window.

### ESC-R encoding genes are highly enriched within *Ec*-BSI sub-populations

We combined the β-lactamase encoding gene data with our WGS based phylogeny to assess the relationship between *E. coli* genetic content and ESC-R mechanisms (**Figure 3**). There was a strong association of phylogroup B2 isolates harboring *bla*_CTX-M-15_ genes as 60.4% of (81/134) *bla*_CTX-M-15_ harboring strains were present in this phylogroup (row-wise Fisher’s exact test adjusted *P*-value = 0.01). Specifically, this association was driven by the presence of *bla*_CTX-M-15_ in 72 STc131 isolates (adj. *P-*value = 0.009), the vast majority of which are part of the previously identified *bla*_CTX-M-15_ carrying *H*30Rx lineage (15). Similarly, there was also enrichment of the second most commonly detected CTX-M encoding gene variant, *bla*_CTX-M-27_, in B2 index isolates (26/29 *bla*_CTX-M-27_ harboring strains; adj. *P-*value < 0.0001) again driven by STc131 isolate carriage (23/29; Adj. *P-*value = 0.0005). Specifically, there were 19/23 *bla*_CTX-M-27_ positive ESC-R-*Ec* belonging to the ST131-C1-M27 clade, which has a strong association with *bla*_CTX-M-27_ carriage (28) as seen in the lower right quadrant of the core gene phylogeny (**Figure 3**). The third most prevalent CTX-M encoding gene variant, *bla*_CTX-M-55_, was enriched within phylogroup A (12 phylogroup A strains out of 26 *bla*_CTX-M-55_ containing index isolates; adj. *P-*value = 0.004) whereas it was only present in one B2 isolate (1/26, Adj. *P-*value < 0.0001). The final major *bla*_CTX-M_ variant in our cohort, *bla*_CTX-M-14_ was carried by a diverse array of genetically heterogenous strains (**Figure 3**). In contrast to the *bla*_CTX-M_ encoding genes in which there were several large clusters of genetically adjacent strains encoding the same CTX-M variant, carriage of CMY encoding genes was sporadically observed across phylogroups. Similar to the phylogroup trend and phylogeny analyses, we did not observe any statistically significant association between ESC-R mediating enzyme variants and infection windows (χ^2^ adj. *P*-value > 0.05).

### Accessory genomes are distinct between B2 and non-B2 phylogroups

We further explored genomic differences within ESC-R-*Ec* by comparing accessory genome content (*i.e.,* defined as genes carried in >5% and <95% of the total ESC-R-*Ec* population) across the full cohort. There was a total of 25,631 genes in the pan genome with 6,060 accessory genes. Through a cluster analysis using the Jaccard distance of the accessory genome between samples, we observed a strong association between phylogroup, clonal complex, and the accessory genome (**Figure 4**). A neighbor joining (NJ) tree constructed from a Jaccard distance matrix shows a clear delineation of B2 and non-B2 isolates that split at the root of the tree (**Figure 4**). Using a network graph analysis of the accessory genome (see methods) (Fig. S4), there was separation of B2 and non-B2 isolates into ten clusters highly correlating to STc (**Figure 4**). Indeed, each of the clonal complexes with ≥5 isolates had at least one statistically significant association (Adj. *P*-value <0.05) with an accessory genome Markov Cluster (MCL) group (Fig. S5). Furthermore, the MCL clustering algorithm delineated the sub-clade structure of STc131 where each of the ST131-*H*30Rx subclades C1 (Cluster 2) and C2 (Cluster 3) were separated by accessory genome (**Figure 4**). ST131-*H*41-O16 subclade A (Cluster 9) was found exclusively in one MCL group (**Figure 4**) demonstrating unique accessory genome between ST131 C1/C2 and A subclades. A complimentary principal component analysis demonstrated clustering of accessory content by phylogroup (Fig. S6A), STc (Fig. S6B), MCL group (Fig. S6C), and ESBL gene (Fig. S6D) further highlighting the separation of B2 and non-B2 groups by accessory genome content. The emergent ST1193 lineage, which made up all STc14 (n=15) strains, made up a separate MCL cluster (Cluster 8) within B2 isolates, however, there were no clear associations with a particular *bla*_CTX-M_ variant. These analyses indicate potential differential pathways to ESC-R development dependent on phylogroup background.

**Figure 4.**
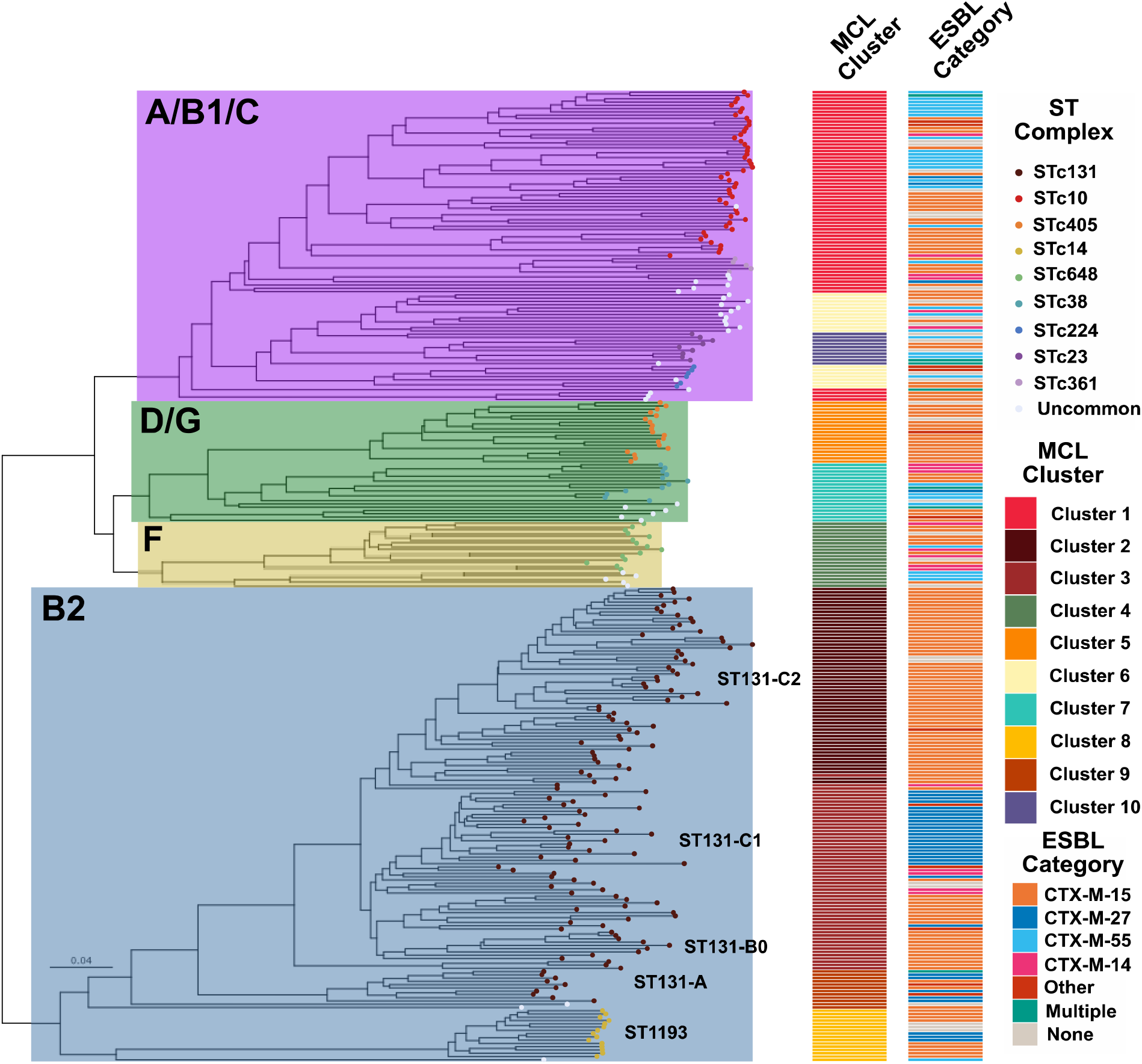
Accessory Genome Clustering by ST131 and non-ST131 ESC-R Isolates. Jaccard distance matrix inferred NJ tree of accessory genome content with metadata indicated in legend. Phylogroups and chosen STs/clades are highlighted within tree and labelled accordingly. MCL = Markov Clustering group; STc = Sequence Type Clonal Complex

### Significant copy number amplifications of enriched CTX-M encoding genes occur within phylogroups

Given the growing appreciation of β-lactamase gene amplification in contributing to AMR phenotypes (29–31), we sought to characterize gene copy number variation of ESC-R genes to determine whether ESC-R gene amplification correlated with our previously identified gene enrichment associations. **Table 3** provides copy number estimates of *bla*_CTX-M_ variants stratified by the three most common phylogroups. Tables S4 and S5 provide copy numbers of other β-lactamase encoding genes and overall estimated copy numbers of AMR genes respectively. For the 268 *bla*_CTX-M_ genes, the median copy number variant (CNV) was 1.2× (Interquartile Range [IQR] = 1.3) with a range of 0.4 to 12.9. The median CNV across CTX-M variants was similar with *bla*_CTX-M-15_ (n=161, CNV=1.1×; IQR =1.2), *bla*_CTX-M-27_ (n=37, CNV=1.4×; IQR =0.8), *bla*_CTX-M-55_ (n=37, CNV=1.5×; IQR =1.4), and *bla*_CTX-M-14_ (n=37, CNV=1.4×; IQR =2.2) having statistically significant greater than 1 copy (one-sample Wilcoxon signed rank adj. *P*-value <0.01).

We and others have previously identified that copy number increases in narrow-spectrum and extended-spectrum β-lactamase encoding genes contribute to the development of carbapenem resistance (32, 33). Therefore, we stratified our ESC-R-*Ec* population by carbapenem susceptibility (*i.e.,* carbapenem non-susceptible vs. susceptible) to determine if there were statistically significant amplifications in carbapenem-susceptible ESC-R-*Ec* isolates. The carbapenem non-susceptible ESC-R-*Ec* had greater estimated copy numbers of *bla*_CTX-M_ (median CNV = 2.7×; IQR = 4.2) compared to carbapenem susceptible ESC-R-*Ec* (median CNV = 1.1×; IQR = 1.0; two-sample Wilcoxon rank sum test *P*-value = 3e-7). Thus, we further characterized *bla*_CTX-M_ gene amplification across phylogroup solely within ESC-R-*Ec* that are carbapenem susceptible (**Figure 5A**). We found statistically significant amplifications for *bla*_CTX_-_M-15_ and *bla*_CTX-M-27_ in phylogroup B2 isolates and *bla*_CTX-M-55_ in phylogroup A isolates, which aligned with the enrichment analysis. These statistically significant trends persisted specifically within STc131 for *bla*_CTX-M-15_/*bla*_CTX-M-27_ and STc10 for *bla*_CTX-M-55_ (Fig. S7). To assess whether *bla*_CTX-M_ amplifications occurred as an adaptive response, we sought to identify isolates from the same patient that were the same clonal complex and had the same *bla*_CTX-M_ variants, *i.e.,* were likely clonal, recurrent infections (**Figure 5B**). We identified 16 pairs from 14 patients in which 56% (9/16) second ESC-R *Ec*-BSI isolates were carbapenem non-susceptible following their antecedent carbapenem susceptible isolate. We found that there was a statistically significant increase (Wilcoxon Signed Rank Test *P*-value=0.03) in *bla*_CTX-M_ copy numbers from the first isolate (median CNV=1.02; IQR=0.86) to the second isolate (median CNV=1.80; IQR=3.32). Notably, these *bla*_CTX-M_ amplifications occurring within sequential isolates were observed for *bla*_CTX-M-15_/*bla*_CTX-M-27_ and *bla*_CTX-M-55_ in STc131 and STc10 respectively (**Figure 5B**). To gain insights into how β-lactamase gene amplification affected AMR phenotypes, we characterized the minimum inhibitory concentration (MIC) distribution for recurrent vs. non-recurrent (*i.e.*, first isolate) populations for piperacillin-tazobactam (TZP), ceftazidime (CAZ), cefepime (FEP), ertapenem (ETP), and meropenem (MEM) antimicrobial susceptibility testing (Fig. S8A-D). For all tested β-lactams, the MIC90 was higher for the recurrent relative to the non-recurrent isolates (Fig. S8). Thus, there was a general shift in MIC distributions which correlated with recurrent isolates that had increased *bla*_CTX-M_ gene copy numbers.

**Figure 5.**
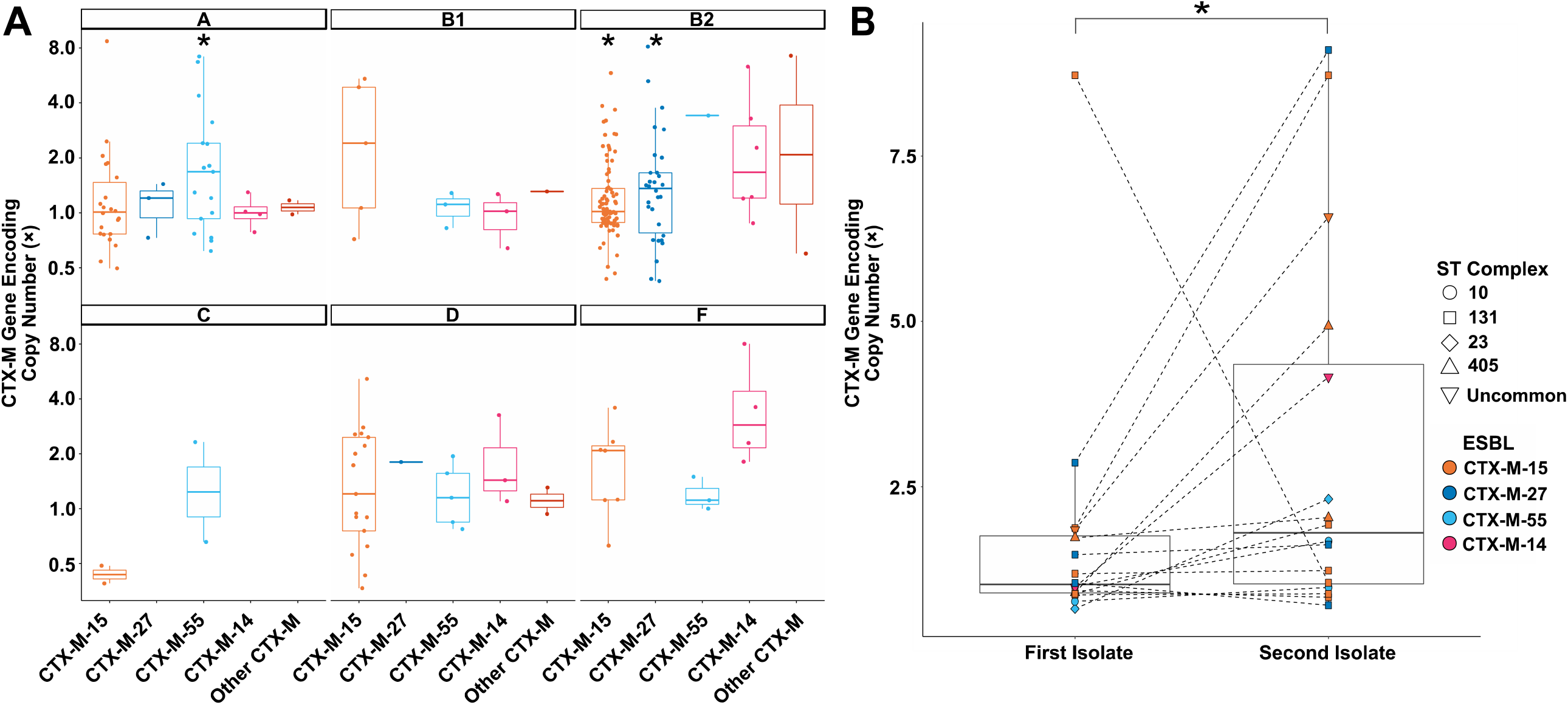
*Bla*_CTX-M_ Amplification Occurring within ESC-R-*Ec* Isolates. **(A)** Copy number estimates of *bla*_CTX-M_ variants stratified by phylogroup indicated in facetted plots that are ESC-R and carbapenem susceptible. One-sample Wilcoxon rank sum test for amplification (*i.e.*, copy number > 1) performed across each *bla*_CTX-M_ variant with * indicating *P-*value<0.05 **(B)** Estimated copy number of *bla*_CTX-M_ variants of paired ESC-R serial isolates (selection criteria as outlined in the text). Shapes and colors indicate clonal complex and CTX-M variant as noted in the legend. Dotted lines connect paired isolates. Paired Wilcoxon-signed rank test was statistically significant with * indicating *P-*value<0.05.

### F-type plasmids are primary vectors of *bla*_CTX-M-55_ in non-STc131 strains

Previous studies have focused largely on ST131 subclade associations with *bla*_CTX-M-15_ and *bla*_CTX-M-27_ (15, 28) including our characterization of *bla*_CTX-M-15_ and *bla*_OXA-1_ co-carriage on similar, multireplicon F-type plasmids contributing to carbapenem resistance (32, 33). Given the rarity of *bla*_CTX-M-55_ molecular epidemiology data from North America (34), we focused on *bla*_CTX-M-55_ inasmuch as it was highly prevalent throughout our sampling frame. We used a long-read approach to complete assemblies of nine *bla*_CTX-M-55_ positive strains and found that most (n=7) carried *bla*_CTX-M-55_ in a highly conserved IS*Ecp1* context on ∼100kb IncFIB/IncFIC plasmids (**Figure 6A**). Whereas the cargo gene region was relatively variable, the F-type plasmid regions responsible for transfer, replication, and stability were highly conserved (blastn ID/coverage ≥99%). For the full cohort of *bla*_CTX-M-55_ isolates (n=38), 31 isolates had IncFIB/IncFIC amplicon hits from PlasmidFinder (>80% ID;>80% COV). Short-read pileup analysis to pMB8960_1 (*i.e.,* the most conserved IncFIB/IncFIC *bla*_CTX-M-55_ positive isolate), showed that 30/31 isolates had >80% breadth of coverage and >100× coverage depth for pMB8960_1.

**Figure 6.**
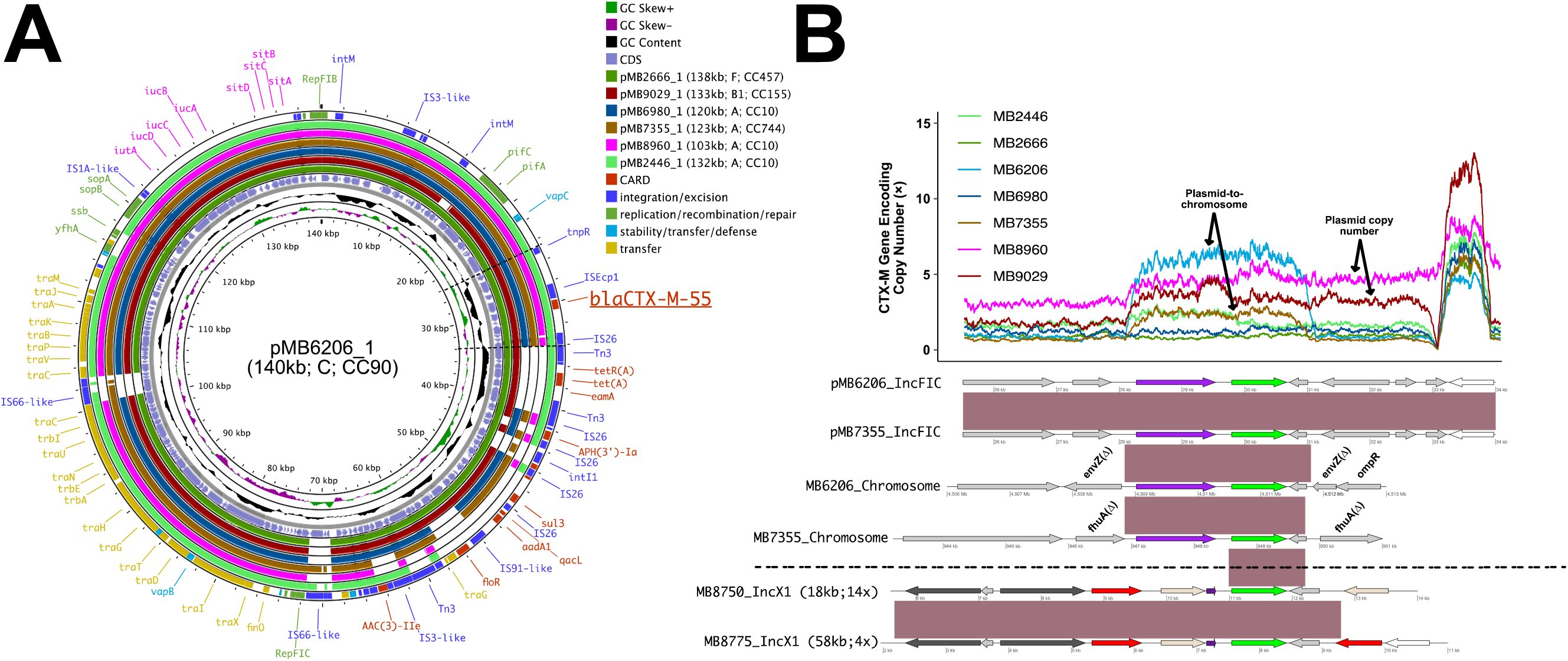
Characterization of mobile genetic elements associated with transmission and amplification of *bla*_CTX-M-55_ within non-B2 isolates. (A) BRIGs output of similar IncFIB/IncFIC plasmids with conserved carriage of IS*Ecp1*-*bla*_CTX-M-55_ carriage (Underlined) in 4 different non-B2 phylogroups. Inner ring indicates GC content/skew with coding DNA sequences in reverse/forward orientation. Next seven rings indicate blastn identities to pMB6206_1 as indicated in legend. Outer ring includes plasmid, mobile genetic element, and antimicrobial resistance gene content as identified by the Comprehensive Antimicrobial Resistance Database (CARD). (B) Blast nucleotide and gene amplification comparisons of differential *bla*_CTX-M-55_ vectors responsible for plasmid transmission as well as potential vertical transmission via chromosomal insertion. Normalized coverage depth of conserved ∼10 kb region that harbors IS*Ecp1*-*bla*_CTX-M-55_ (∼10kb) as demarcated by dotted black lines in (A). Inferred plasmid copy number vs. plasmid-to-chromosome events as indicated by tracking target site duplications of IS*Ecp1*-*bla*_CTX-M-55_ on complete assemblies as well as mean coverage depths of full-length plasmid. Coding sequences within blastn comparisons are identified as followed: IS*Ecp1* (purple), *bla*_CTX-M-55_ (green), AMR genes (red), IS*26* (white), IS*26*-v1 (beige), other MGE (black), other CDS (gray). Black horizontal line demarcates IncFIB/IncFIC positive isolates from high copy number IncX1 plasmids harboring *bla*_CTX-M-55_ from ST167 isolates. Parentheticals following plasmid label indicate plasmid length and copy number. CDS with striped, purple label indicates IS*Ecp1* truncated by IS*26*-v1 upstream of *bla*_CTX-M-55._

Our long read analyses allowed for identification of *bla*_CTX-M-55_ amplification arising from plasmid copy number variation (MB8960 and MB9029) as well as plasmid-to-chromosome transfer (*i.e.,* segmental duplication) from two isolates (MB6206 and MB7355) as evidenced by tracking target site duplications (TSDs) and plasmid content from IS*Ecp1*-*bla*_CTX-M-55_ (**Figure 6B**). This included an isolate in which IS*Ecp1*-*bla*_CTX-M-55_ disrupts the *envZ* gene, which is part of a two-component regulatory system (*i.e.,* EnvZ-OmpR) that regulates expression of outer membrane porin genes (35). A carbapenem susceptible ESC-R-*Ec* isolate (MB7355) was identified with two independent insertions within the chromosome likely originating from the IncFIB/IncFIC plasmid as evidenced through TSD tracking (**Figure 6B;** one insertion into *fhu*A gene demonstrated). Another vector of *bla*_CTX-M-55_ was identified in two ST167 isolates in a substantially differing context with *bla*_CTX-M-55_ upstream of IS*26*-v1 on smaller IncX1 plasmids (**Figure 6B**). The IS*26*-v1 upstream of *bla*_CTX-M-55_ had truncated IS*Ecp1* (**Figure 6B;** purple striped CDS) and were within unique antimicrobial resistance gene transposons. These analyses together show that *bla*_CTX-M-55_ is a major contributor to the ESC-R-*Ec* phenotype in our cohort with predominantly IncFIB/IncFIC carriage.

## DISCUSSION

The past twenty years in the US have seen a consistent rise in ESC-R-*Ec* causing infection, primarily due to ESBL enzymes (2–4). While the high prevalence of ST131 observed in the US is concerning, it is not clear if these pathogens are responsible for the increased frequency of ESC-R-*Ec* (2). We sought to leverage our longitudinal collection of bloodstream isolates to address the recent temporal dynamics of *E. coli* at our tertiary care cancer center. Although we observed that STc131 strains accounted for nearly 50% of infecting isolates, the STc131 prevalence rates remained fairly stable over time with multiple MDR clones accounting for peaks in infection rates observed during our study period.

We chose to focus on BSIs because of the clear distinction between colonization and infection relative to other potential sample sites. Our large number of fully sequenced isolates and extended study timeframe provide critical information regarding contemporary molecular epidemiology of invasive ESC-R-*Ec* infections in a major US city. Although our finding of ∼35% *Ec*-BSIs being ESC-R is high relative to other studies from the US (4), it is consistent with the ∼40% proportion of ESC-R in *Ec*-BSI occurring in solid-organ transplant recipients (36). Whereas recent longitudinal ESC-R-*Ec* epidemiological studies from the US are lacking, we can compare our findings to those recently published from Europe (16–21). A similar theme has emerged from these investigations where for several years following introduction into the population, STc131 strains dramatically increase before becoming a stable proportion of overall *E. coli* infections (16–21). Given that STc131 had been identified as the dominant antimicrobial resistant *E. coli* STc in the US for at least 10 years prior to the start of our sampling (27, 37), our finding of a relatively stable contribution of STc131 is consistent with other studies (16–21) and suggest that, at least for our population, STc131 may have reached a stable level in the broader ESC-R-*Ec* population.

A key strength of our study was our ability to investigate the molecular epidemiology underlying the temporal variability of ESC-R-*Ec* prevalence rates. We found that ESC-R-*Ec* rates generally peaked within the last six months of the calendar year and were independent of ESC-S-*Ec* rates. Seasonality of both ESC-R-*Ec* infection and colonization have been previously reported with generally higher rates identified during warmer months (38–40). For example, a recent pooled observational cohort analysis of ESBL positive *E. coli* and *K. pneumoniae* (ESBL-E/K) carriage in the Netherlands found that the highest prevalence occurred in August (OR 1.88, 95% CI 1.02-3.49) and September (OR 2.25, 95% CI 1.30-3.89) compared to January (39). In Houston, the highest temperatures of the year are generally observed between June and September (https://www.weather.gov/hgx/climate_iah_normals_summary), which might facilitate proliferation of *E. coli* strains leading to colonization and subsequent infection at a delayed interval. Consistent with environmental factors being a major driver of the temporal variability, we observed low genetic relatedness amongst *E. coli* strains during the peak windows of infection rather than closely related clones. On a few occasions, we identified strains from distinct patients that were closely related (*i.e.,* < 10 SNPs) consistent with hospital acquisition, but most strains were genetically diverse suggesting a community source. Given the recent success of wastewater surveillance in identifying SARS-CoV-2 and influenza transmission dynamics (41, 42), we envision that a similar approach could be used to improve understanding of how community prevalence of ESC-R pathogens impacts subsequent infections.

In previous works we have identified that non-carbapenemase producing, carbapenem resistant *Enterobacterales* (non-CP-CRE) have increased copy numbers of genes encoding ESC-R mediating enzymes such as *bla*_CTX-M_ variants (32, 33). This study expands on our previous copy-number assessment of β-lactamase encoding genes mediating the ESC-R phenotype in carbapenem-susceptible *E. coli* strains. This allowed us to find that carbapenem-susceptible ESC-R-*Ec* strains have lower *bla*_CTX-M_ copy numbers relative to carbapenem-resistant ESC-R-*Ec* strains consistent with an important role of *bla*_CTX-M_ amplification in the transition from ESC-R to the non-CP-CRE phenotype. Moreover, via an analysis of paired isolates from the same patients, we found significant *bla*_CTX-M_ amplifications suggesting that such amplifications are likely an adaptive mechanism to antimicrobial exposure in the host as has been observed by our group and others in the laboratory environment (32, 33, 43). Together with recently published data from ESBL-positive *E. coli* strains isolated in St. Louis (34), our long-read analyses demonstrate the diverse mechanisms by which *E. coli* can increase *bla*_CTX-M_ encoding gene copy numbers including augmented plasmid copy numbers, multiple chromosomal copies in distinct locations, and *in situ* amplifications mediated by insertion sequences such as IS*Ecp1* and IS*26*. Although gene amplifications have long been thought to confer a major fitness cost and thus be transient in response to antimicrobials, incorporation of *bla*_CTX-M_ encoding genes into particular chromosomal locations has been found to have a low fitness cost, allowing for stable amplifications across many generations in the absence of antimicrobial pressure (44, 45). We hypothesize that the high rates of recurrence observed amongst ESC-R-*Ec* relative to ESC-S-*Ec* strains might, in part, be due to *bla*_CTX-M_ amplifications occurring amongst colonizing strains in the gastrointestinal tract which would allow for their persistence during carbapenem treatment and subsequent re-infection. Such an amplification of *bla*_CTX-M-14_ was recently described in an *E. coli* strain recurrently isolated from a patient’s urine over many months (34).

Furthermore, we were able to identify IncFIB/IncFIC plasmids as the primary vector of IS*Ecp1*-*bla*_CTX-M-55_ carriage in our cohort where amplification was occurring through both increase in plasmid copy numbers as well as plasmid-to-chromosome horizontal gene transfer. Although we did not identify a ‘clonal expansion’ of chromosomal IS*Ecp1*-*bla*_CTX-M-55_, the independent transfer of these vectors into two different phylogroup chromosomal backgrounds suggest the potential for stable chromosomal propagation. A recent study in China found a temporal trend with increasing chromosomal *bla*_CTX-M-55_ albeit without clarity to whether this was occurring within clonal populations (46). While little is known regarding *bla*_CTX-M-55_ prevalence in North America (34), the enrichment of this *bla*_CTX-M_ variant in the second most commonly detected phylogroup/STc in our cohort suggest these IncFIB/IncFIC::IS*Ecp1*-*bla*_CTX-M-55_ plasmids are a significant contributor to ESC-R-*Ec* in our region.

Although our study has many strengths, there are important limitations. First, this was a single-center investigation at a hospital serving an immunocompromised population and thus external generalizability is unclear. This weakness is somewhat mitigated by our center serving a geographically dispersed group of patients and because the impact of ESC-R-*Ec* is particularly severe amongst immunocompromised persons. Moreover, our finding of temporal variability amongst *E. coli* infections has been observed amongst broader populations such as a recent nationwide report from Israel (47). Second, the diverse nature of observed clonal complexes and subclones meant that there were often small numbers of non-STc131 clonal complexes during specific infection windows. Larger sampling from a more geographically dispersed area is needed to address these issues. A final weakness is that the beginning of the COVID epidemic likely led to a major disruption in infection epidemiology at our institution given the nearly 50% decrease in admissions observed in the spring and summer of 2020. Whereas both the absolute numbers of and proportion of ESC-R-*Ec* BSIs had been steadily increasing from 2017 through 2019, both significantly decreased during 2020. The reasons for these findings are not clear, but they stand in contrast to preliminary data released by the CDC indicated an increase in ESBL Enterobacterales in 2020 nationwide (48).

In summary, we present a comprehensive whole genome-based analyses of ESC-R-*Ec* from the US analyzing recently collected strains. These data show that STc131 strains have reached a stable proportion of our ESC-R-*Ec* population. Additionally, we provide molecular insights into the temporal variability of ESC-R-*Ec* prevalence which is being increasingly appreciated but is not well understood mechanistically (16–21). The genetically diverse nature of ESC-R-*Ec* strains identified in our study indicate that efforts to mitigate serious ESC-R-*Ec* infections likely will need to address broad environmental reservoirs such as through vaccination and highlight the need to better understand ESC-R-*Ec* transmission and temporal dynamics outside of the hospital setting.

## METHODS

### Study design and sample collection

*Escherichia coli* bloodstream infections (*Ec*-BSI) occurred from March 1^st^, 2016, to December 31^st^, 2020, at MD Anderson Cancer Center (MDACC) in Houston, TX with March 1^st^, 2016, chosen given this was the date of implementation of the Epic Electronic Health Record system. We defined an index *Ec*-BSI isolate as first occurrence of a positive blood culture and a recurrent *Ec*-BSI isolate as a positive blood culture that occurred at least 14 days from a previous *Ec*-BSI. Antibiotic susceptibility testing of all *Ec*-BSI isolates was performed by the MDACC microbiology laboratory using the Accelerate PhenoTest™ BC (Accelerate Diagnostics), ETEST® (bioMérieux), and VITEK® 2 (bioMérieux). *Ec*-BSI Isolates were considered as extended-spectrum cephalosporin resistant (ESC-R) if they had a ceftriaxone (CRO) minimum inhibitory concentration (MIC) ≥ 4 μg/mL and/or a predicted ESBL phenotype.

### Genomic sequencing

ESC-R-*Ec* isolates were sequenced by Illumina NovaSeq 6000 or Illumina MiSeq as previously described (33). There were 297 ESC-R isolates that had paired-end 150 base pair reads that passed quality control assessed using fastqc-v0.11.9. We performed long-read sequencing on an nine isolates using the Oxford Nanopore Technologies GridION platform as described previously (33).

### Genome analysis and molecular typing

Quality assessed, paired-end 150 bp reads were assembled via SPAdes-v3.15.2 (49) with default parameters and the inclusion of the isolate option. Quality assessment of output SPAdes assembly contigs was performed via QUAST-v5.2.0 (50) to determine N50 and contiguity. Assemblies with (a) genomes < 4 Mb or > 6 Mb; or (b) Contigs >500 were excluded from analysis. The EzClermont i*n silico* phylotyping tool was utilized with the 2013 Clermont PCR typing method (Waters, N EzClermont Github: https://github.com/nickp60/EzClermont). *E. coli* multilocus sequence typing (MLST) using the Achtman definition was performed with the mlst-v2.19.0 *in silico* contig scanning tool (Seemann T mlst GitHub: https://github.com/tseemann/mlst), which features the PUBMLST database (51). The Copy Number Variant quantifICation Tool (convict-v1.0) was used to identify antimicrobial resistance (AMR) genes and estimate gene copy numbers as described previously (Shropshire, W convict GitHub: https://github.com/wshropshire/convict) (33). Briefly, short-read data is used as input to KmerResistance-v2.2.0 (52, 53) to identify potential AMR genes from the resfinder database (Accessed 2022-10-01). Through a coverage depth ratio of AMR gene to housekeeping gene that accounts for variance in coverage depths, convict estimates gene copy numbers. The high-performance computing (HPC) cluster, Seadragon, that is hosted through MDACC was used to perform genomic analyses.

### Pan genome analysis

Draft and complete genomes were annotated using the Prokka-v1.14.6 (54) using default parameters. Annotated gff files from prokka were used as input for pangenome analysis using Roary-v3.13.0 (55) using a protein family ID threshold of 95%. A core gene alignment was completed using Mafft-v7.508 with the Roary pipeline using a core gene threshold of 99%. This core gene alignment file was used as input to IQTree2-v2.2.0-beta to create a core gene inferred maximum likelihood phylogeny. ModelFinder was utilized to determine most appropriate DNA model with the best fit model: UNREST+FO+I+G4 chosen according to BIC. A final bootstrap consensus tree was created using 1000 SH-like aLRT replicates in addition to 1000 Ultrafast Bootstrap support values. A core genome alignment was performed on the top 12 sequence types observed in our cohort using Snippy-v4.6.0 using default variant calling parameters. The following were the chosen references for alignment: (ST131: CP049085.2; ST405: CP103633.1; ST1193: MB11100; ST648: CP103657.1; ST744: MB7355; ST10: CP049081.1; ST167: CP103540.1; ST38: CP103559.1; ST617: CP103645.1; ST410: CP103533.1; ST224: CP103609.1; ST361: CP103704.1). A core genome alignment was used as input for Gubbins (56) to create a recombination-free, maximum likelihood tree using IQTree20v.2.2.0 using the aforementioned parameters. The recombination free fasta output was used to extract pairwise SNP sites (snp-sites-v2.5.1) (57) and SNP distances (snp-dists-v0.8.2) respectively. A pairwise SNP distance cutoff of 10 for window independent analyses of transmission was based on the recombination-free, core-genome SNP differences of eight pairwise comparisons of ST131 recurrent isolates where the median pairwise SNP difference was 5.5 SNPs and the maximum observed was nine SNPs.

The accessory genome was assessed using the gene_presence_absence.csv file output from Roary. A Jaccard distance matrix was created using this binary accessory genome file after removing genes present in <5% and >95% of the total population. This distance matrix was used as input to create a neighbor joining tree using the BIONJ algorithm (58). We performed a principal component analysis of this distance matrix using base R functions. The graph network analysis was accomplished using the same gene_presence_absence.csv input using graphia (59, 60). Edge transformations using a k-nearest neighbors algorithm (n=8) were employed to reduce the number of edges from 44k to 1724. Clustering was performed using the Markov Cluster algorithm (MCL) with an inflation value=1.5 (61). Visualization of data was accomplished using ggplot2-v3.3.5 (62) and ggtree-v.3.3.1 (63).

### Long read analysis

A short-and long-read assembly pipeline (Shropshire, W flye_hybrid_assembly_pipeline GitHub: https://github.com/wshropshire/flye_hybrid_assembly_pipeline) was used to close complete genomes of ONT sequenced data as described previously (33). Incomplete assemblies were re-assembled using Unicycler-v0.5.0 and manually curated for errors using short-and long-read pileups and visualizing with the integrated genome browser (IGV-v2.14.1). Comparison of plasmid vectors with BLAST ring image generator (BLAST) was performed using the Proksee server while querying AMR genes with the Comprehensive Antimicrobial Resistance Database (CARD) (64).

### Statistical analysis

Prevalence rates were plotted across time using ggplot2 with a smoothing curve (*i.e.,* a loess function) to visualize prevalence trends. Time series data was assessed for monotonic trends using the Mann-Kendall Trend test and non-monotonic using the WAVK statistic. Correlations between ESC-R-*Ec* and ESC-S-*Ec* were measured with a Pearson’s product moment correlation test. We used χ^2^ test (Fisher’s exact when observations <5) to determine statistical differences in proportions across phylogroup/clonal complexes. A Kruskal-Wallis global test to determine differences in gene copy numbers across groups. Pairwise and one-way Wilcoxon Rank sum tests were to determine group level and single mean differences respectively with a Bonferroni correction for multiple comparisons. All statistics was computed on R (4.0.4) and α=0.05 parameter was used across all tests.

### Data Availability

WGS data sequenced during this study period was submitted to NCBI and can be accessed within BioProject PRJNA924946. WGS data from previous studies can be accessed from BioProject PRJNA603908 and PRJNA836696 (33, 65). All R Scripts can be made available through request of the corresponding author.

## Acknowledgements

We would like to thank the MDACC clinical microbiology lab for all their hard work in identifying, handling, and transferring these pathogenic strains of interest to us for our research projects. Core grant CA016672(ATGC) and NIH 1S10OD024977-01 grant provide funding for the Advanced Technology Genomics Core (ATGC) sequencing facility at MD Anderson Cancer Center. WCS is supported through the National Institute of Allergy and Infectious Diseases (NIAID) T32 AI141349 Training Program in Antimicrobial Resistance. Support for this study was also provided by the NIAID R21AI151536 and P01AI152999 for S.A.S. The research in the A.K. laboratory is supported by NIGMS 1R01GM133904-01 and the Welch Foundation Research Grant AU-1998-20190330. The authors acknowledge the support of the High-Performance Computing for research facility at the University of Texas MD Anderson Cancer Center for providing computational resources that have contributed to the research results reported in this paper.

