## Supplemental Figures for "Temporal Dynamics of Genetically Heterogeneous Extended-Spectrum Cephalosporin Resistant *Escherichia coli* Bloodstream Infections"

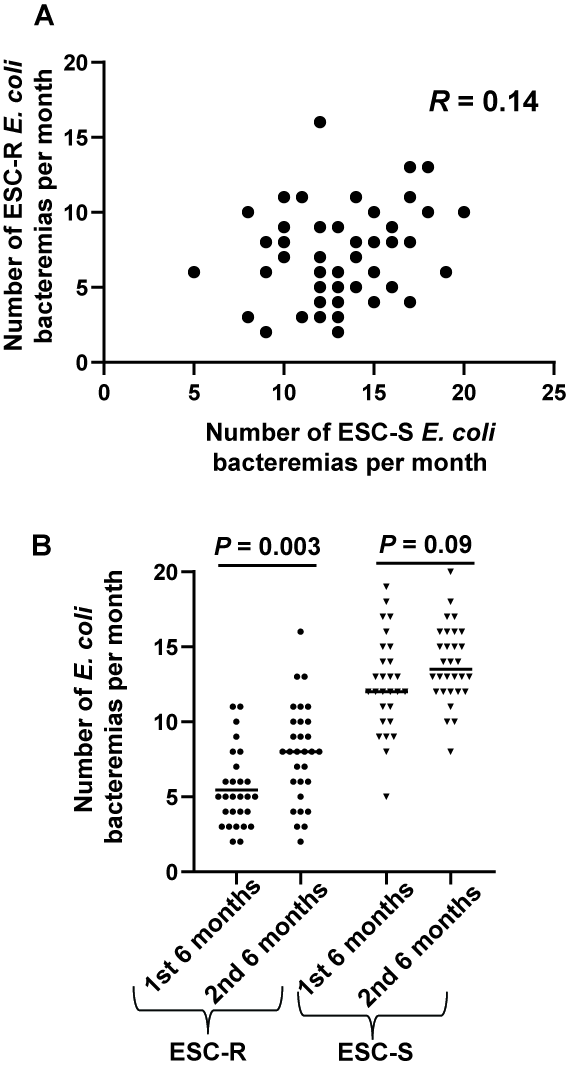

**Fig S1. (A) Correlation plot of extended spectrum cephalosporin-resistant (ESC-R) and susceptible (ESC-S) *E. coli* BSIs per month.** Pearson correlation coefficient *P*-value = 0.32. **(B) Stratification of ESC-R vs. ESC-S *E. coli* BSIs by 6-month time frames.** A statistically significant higher frequency of ESC-R E. coli BSIs was observed per month in 2^nd^ half of year vs 1^st^ half of year (Student’s t-test *P*-value = 0.003).

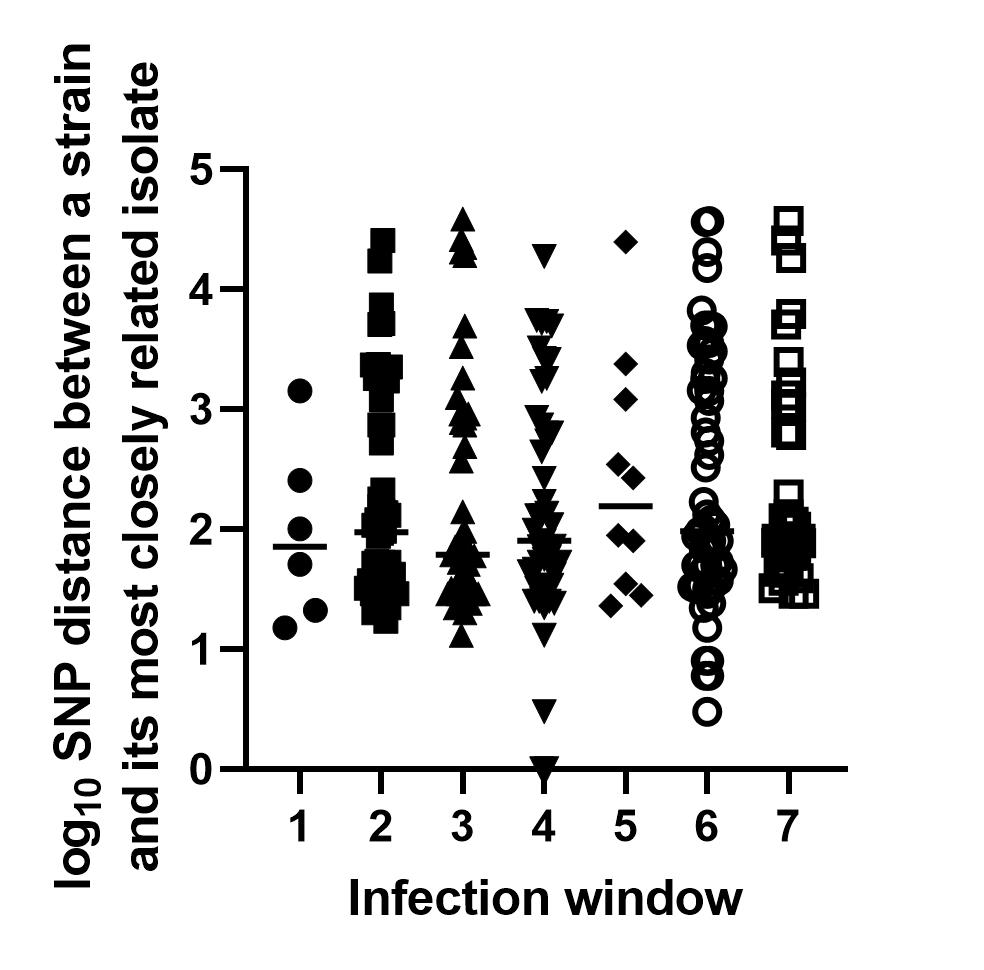

**Fig S2. Nearest Neighbor SNP Distance Across Infection Windows.** Shown are individual single nucleotide polymorphism (SNP) distance for an individual, index ESC-R *E. coli* isolates to its nearest neighbor stratified by infection window. Horizontal bars indicate median values.

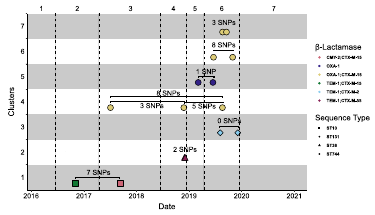

**Fig S3. Time Window Independent Core Genome Pairwise SNP Distance Analysis.** Choosing a pairwise SNP difference cutoff of 10 based on the maximum pairwise distance observed amongst recurrent ST131 isolates, we identified 7 clusters (y-axis) of highly related isolates. Time windows are indicated through vertical dotted black lines. Shape indicates sequence type and color indicates β-lactamase composition as noted in legend.

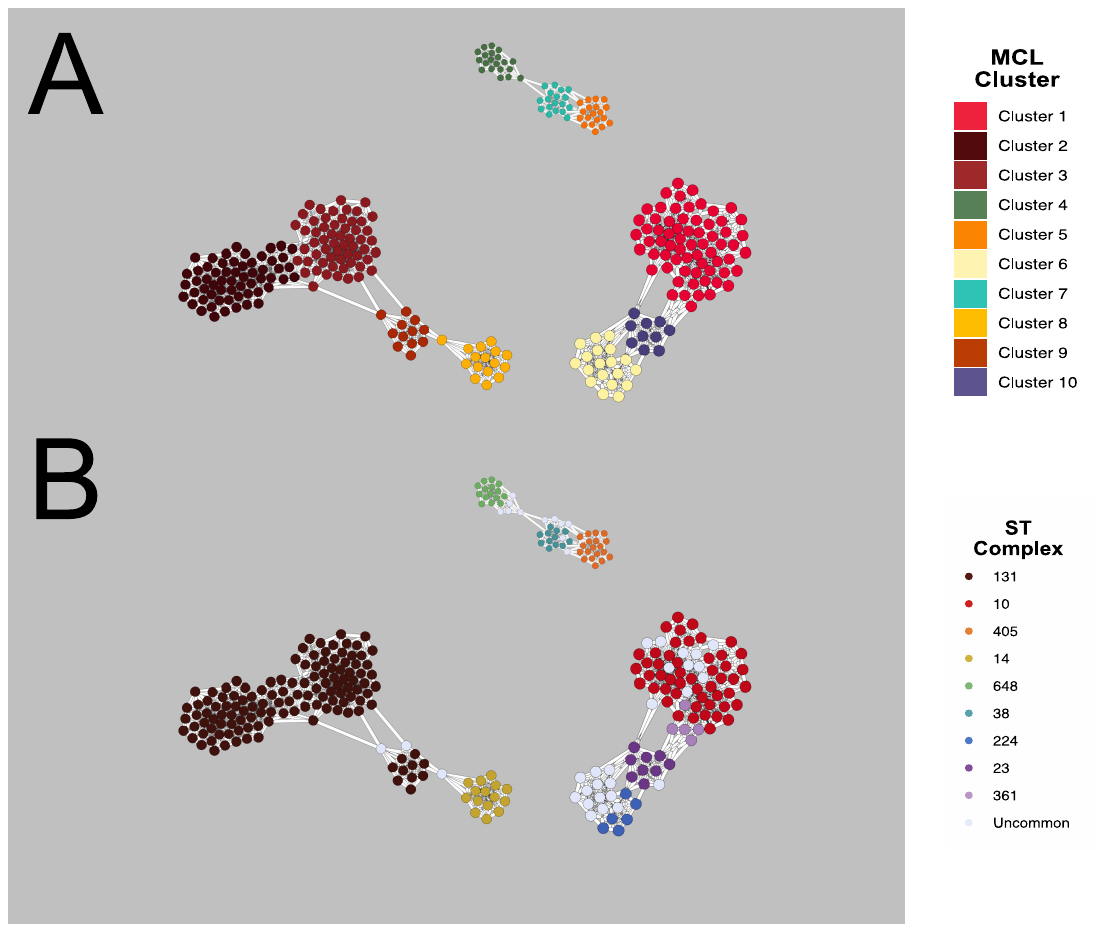

**Fig S4. Network Graph of Accessory Genome.** Graph outlined by **(A)** MCL cluster (n=10) and **(B)** STcomplex. MCL Cluster assignment largely delineates isolates by ST complex, while separating sub-clade structures of ST131, namely C1/C2 from subclade A.

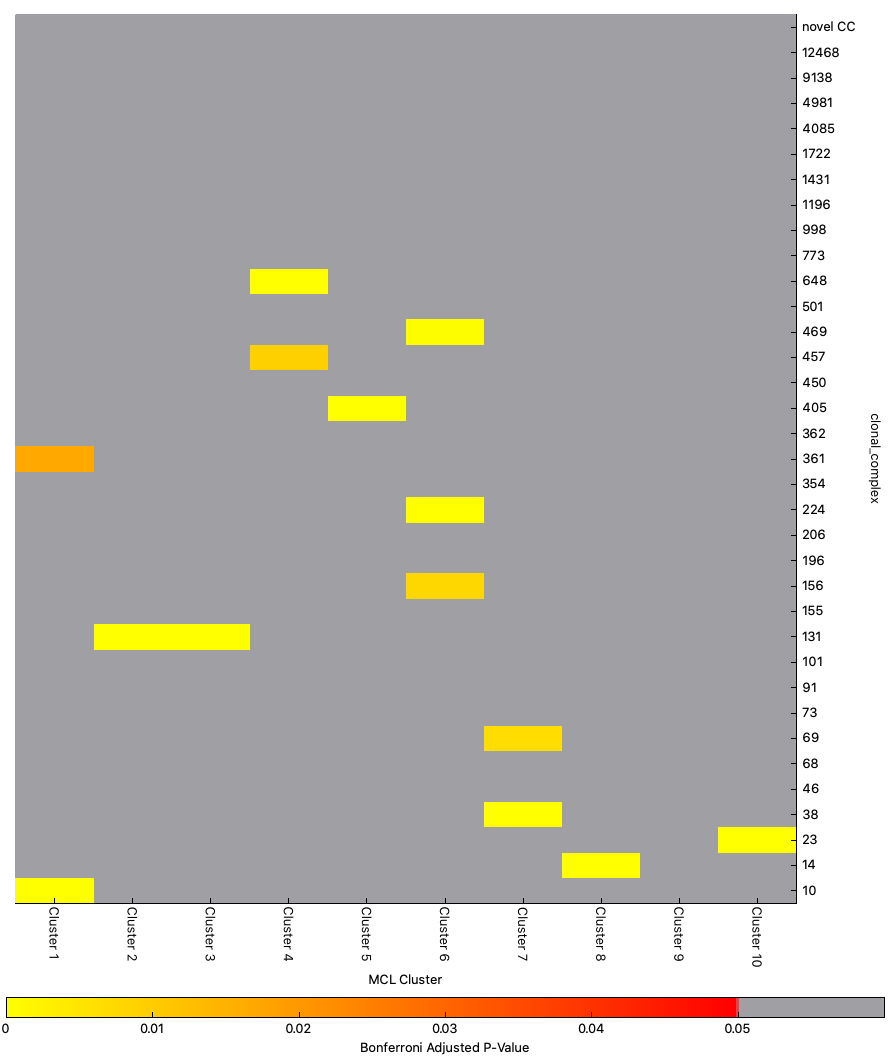

**Fig. S5. Heatmap of Statistically significant Fisher’s Exact Test comparisons of MCL Cluster and ST/ST Complex.** Sequence Type or ST complex is shown on Y-axis with Bonferroni Adj. *P-*value <0.05 indicated within each MCL Cluster on the X-axis. These adj. *P-*values are labelled as a heat-map as presented below the graph.

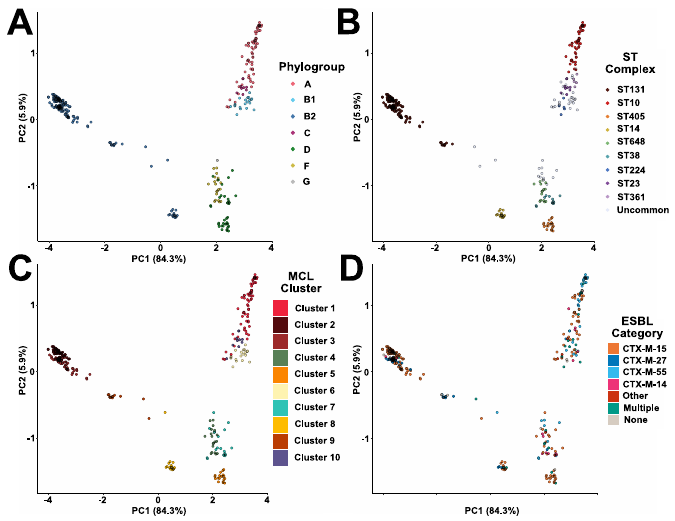

**Fig S6. Principal components, PC1 and PC2, of accessory genome content.** The PC1 and PC2 of accessory genome are plotted with percent variance explained and stratified by **(B)** phylogroup, **(C)** ST complex, **(D)** MCL cluster, **(E)** ESBL status indicated in respective legends.

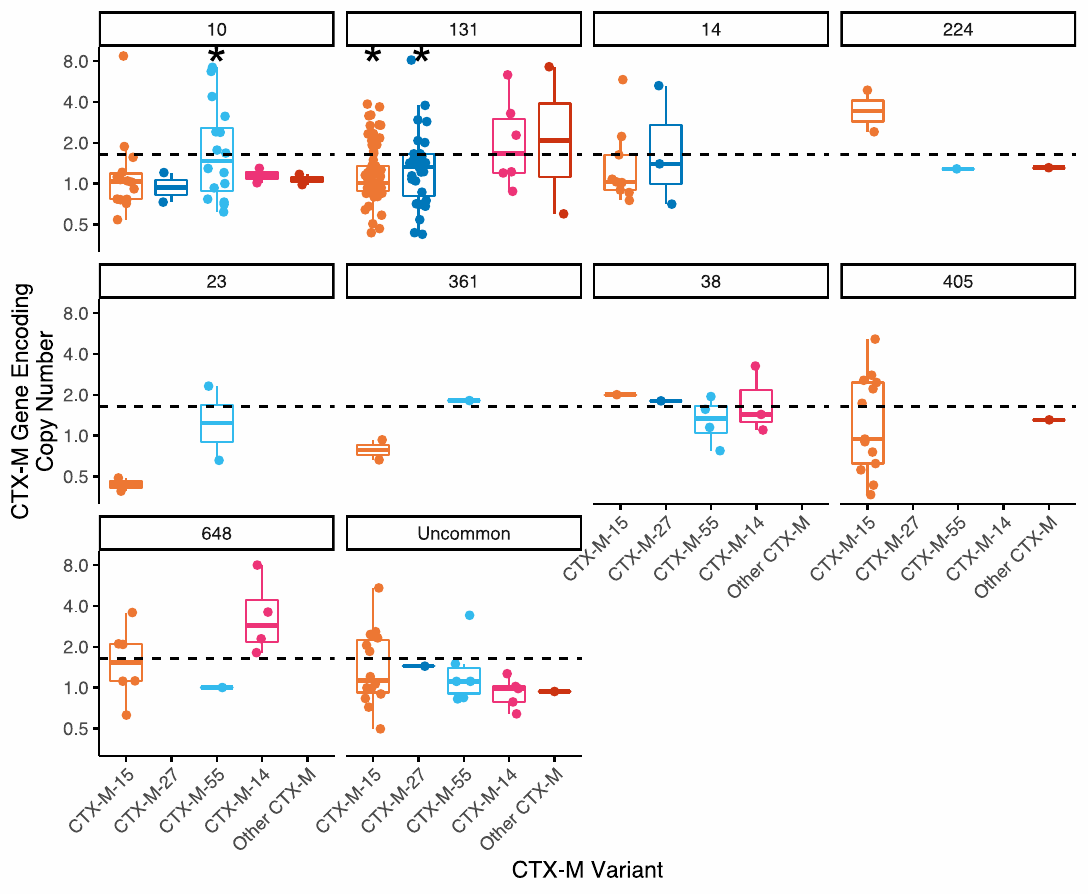

**Fig S7. *Bla*_CTX-M_ Encoding Gene Copy Numbers by ST Complex**. Copy number estimates of *bla*_CTX-M_ variants across ST complex. One-sample Wilcoxon rank sum test performed across each *bla*_CTX-M_ variant with * indicating *P-*value<0.05. Dotted-line indicates mean estimated *bla*_CTX-M_ copy number across ESC-R *Ec*-BSI.

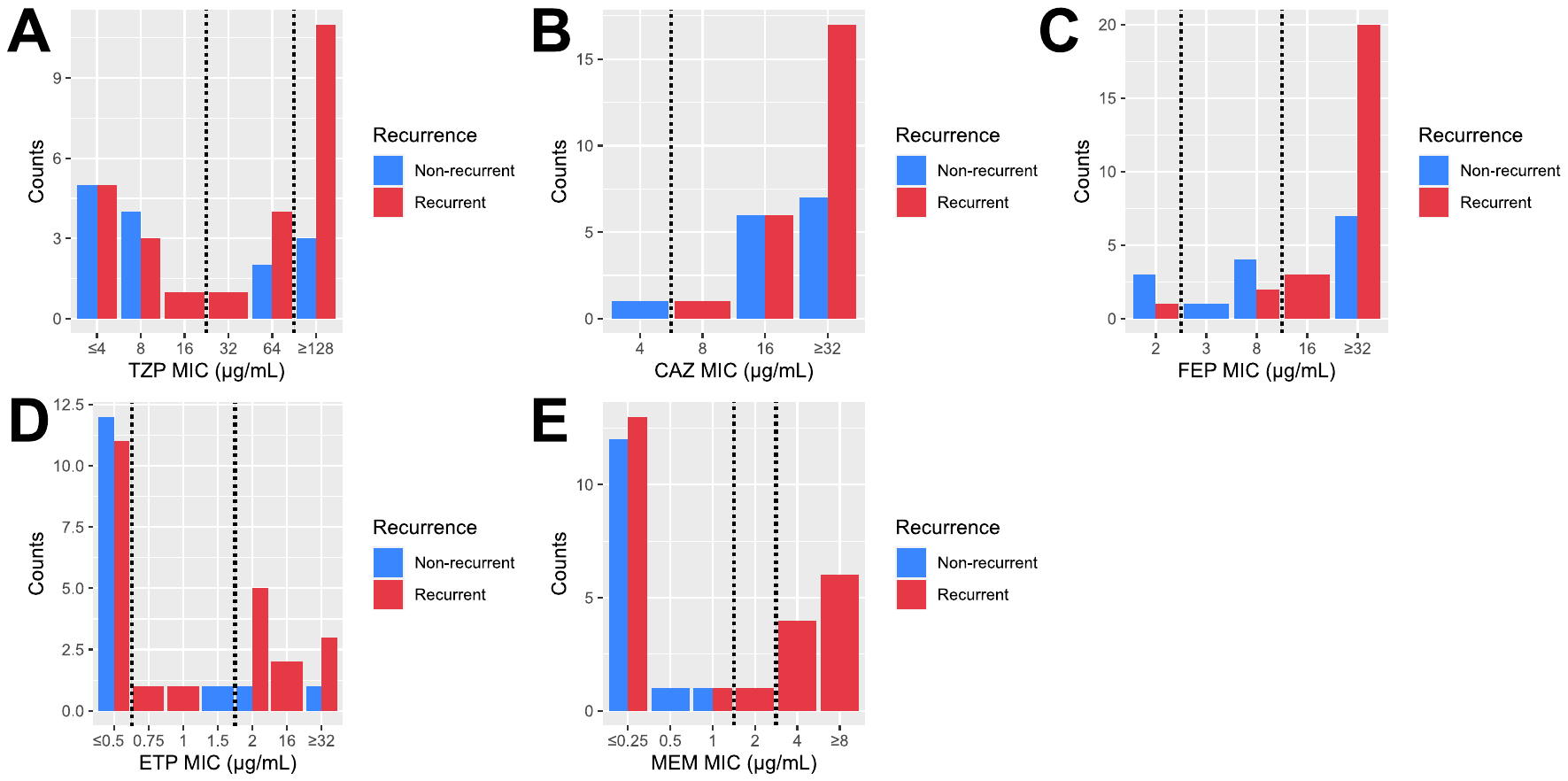

**Fig S8. Minimum Inhibitory Concentration (MIC) Frequency Distribution Across Non-recurrent and Recurrent Isolates. (A)** TZP = piperacillin-tazobactam; **(B)** CAZ = ceftazidime; **(C)** FEP = cefepime; **(D)** ETP = ertapenem; **(E)** MEM = meropenem. Horizontal lines establish MIC cutoffs for susceptible, intermediate, and/or resistant as established by CLSI guidelines (2022).
